# Integrated spatial proteomics of human PDAC uncovers an expanded tumour–immune– stroma spectrum with genomic associations

**DOI:** 10.1101/2025.08.15.670565

**Authors:** Noor Shakfa, Ferris Nowlan, Sibyl Drissler, Tiak Ju Tan, Elizabeth Sunnucks, Edward L. Y. Chen, Cassandra J. Wong, Brendon Seale, Zhen-Yuan Lin, Michelle Chan-Seng-Yue, Amy Zhang, Sabiq Chaudhary, Chengxin Yu, Golnaz Abazari, Michael J. Geuenich, Matthew Watson, Jiaxi Peng, Somaieh Afiuni-Zadeh, Ayelet Borgida, Ricardo Gonzalez, Sheng-Ben Liang, Klaudia Nowak, Miralem Mrkonjic, Anna Dodd, Julie M. Wilson, Kieran R. Campbell, Jennifer L. Gorman, Barbara T. Grünwald, Robert C. Grant, Jennifer J. Knox, Faiyaz Notta, Anne-Claude Gingras, Steven Gallinger, Grainne M. O’Kane, Hartland W. Jackson

**Affiliations:** Lunenfeld-Tanenbaum Research Institute, Mount Sinai Hospital, Toronto, ON, Canada; Department of Molecular Genetics, University of Toronto, Toronto, ON, Canada; Ontario Institute for Cancer Research, Toronto, ON, Canada; Data Sciences Institute, University of Toronto, ON, Canada; Toronto General Hospital, Toronto, ON, Canada; Princess Margaret Cancer Biobank, University Health Network, Toronto, Ontario; Department of Laboratory Medicine & Pathobiology, Toronto, ON, Canada; Wallace McCain Centre for Pancreatic Cancer, Princess Margaret Cancer Centre, University Health Network, Toronto, ON, Canada; Department of Statistical Sciences, University of Toronto, Toronto, ON, Canada; Department of Computer Science, University of Toronto, Toronto, ON, Canada; West German Cancer Center, Universitätsklinikum Essen, Essen, Germany; Department of Urology, University Hospital Essen, Essen, Germany; Department of Medical Biophysics, University of Toronto, Toronto, ON, Canada; Department of Surgery, University of Toronto, Toronto, ON, Canada; Hepatobiliary/Pancreatic Surgical Oncology Program, University Health Network, Toronto, ON; St. Vincent’s University Hospital, School of Medicine, University College Dublin, Dublin, Ireland

**Keywords:** pancreatic cancer, tumour microenvironment, spatial, epithelium, immune, stroma, proteome, transcriptome, genome

## Abstract

Distinctively, pancreatic ductal adenocarcinoma (PDAC) consists of sparse tumour lesions intertwined with extensive desmoplastic stroma. The complexity of tumour–microenvironment interactions within this desmoplasia poses a challenge for accurate tumour profiling and patient stratification, and characterizes a profoundly chemoresistant tumour. Here we mapped the spatial relationships between tumour, stroma, and immune cell compartments delineating tumour and microenvironment types that expand the classical to basal spectrum of human PDAC. We used imaging mass cytometry to profile the *in situ* multi-cellular organization of 81 cell types in resected cases with paired whole genome sequencing. Cell types, functions, and pathway activation were distributed as highly reproducible environments in discrete locations throughout these tumours, which we deep-profiled using laser-capture mass spectrometry. We show that the connections between tumour phenotypes, vascularization, immune response, and stromal biophysical state are reinforced by genomic aberrations, altered by treatment, and associated with patient outcome. Predictive machine-learning models showed that spatial single cell data outperformed genomic or clinical features but integrated multi-omics models provide the best prediction of patient survival with compressed models requiring only 10 non-redundant robust molecular measures associated with the phenotypic spectrum of PDAC. Together, these findings define a phenotypic and molecular framework of PDAC that captures tumour–microenvironment co-dependencies and offers a refined basis for patient stratification and therapeutic targeting.

## Main

With increasing incidence and dismal survival rates, PDAC is a leading cause of cancer death^1^. As most patients present with advanced, inoperable disease, and current conventional therapies offer limited benefit, there is an urgent need for patient stratification and development of targeted therapeutic options^2^.

PDACs contain an intermixed tumour microenvironment (TME) consisting of cancer associated fibroblasts (CAFs), mural and endothelial cells, lymphoid and myeloid populations, and a rich extracellular matrix with stromal content constituting up to 80% of the tumour mass. This desmoplastic stroma is also central to PDAC therapeutic resistance acting as a mediator of chemoresistance and a biophysical barrier to drug delivery and immune infiltration^3^. Histopathology- and transcriptomics-based investigations into specific cell types have identified multiple CAF subtypes, including inflammatory CAFs (iCAF), myofibroblastic CAFs (myCAF), and CD105+ CAFs^4,5^, while other studies have described tissue-level stromal patterns such as activated/normal or reactive/deserted stroma^6,7^. These classification systems have partial overlap despite different methodologies, but are not easily unified^8^. In contrast, transcriptomic studies of cancer-specific expression have identified highly reproducible tumour subtypes called classical (i.e. pancreatic progenitor, or classical A/B) and basal (i.e. quasi-mesenchymal, squamous, basal-like, or basal-like A/B)^6,9–11^. However, 30–45% of tumours are of mixed transcriptomic subtype, with most containing both classical or basal cell types and even dual-expressing (hybrid) cell states simultaneously^10,12,13^. This complexity can confound and limit the study of low cellularity tumour cells, but has led to signatures which can support therapeutic decisions. Basal tumours are generally more aggressive and resistant to standard of care while classical tumours demonstrate less sensitivity to KRAS and panRAS inhibitors (KRASi)^14,15^.

PDAC genomes are characterized by a relatively homogenous landscape of somatic driver mutations, with nearly 90% of tumours harboring activating mutations in *KRAS*, followed by inactivation of *TP53* (∼80%), *CDKN2A* (∼60%), and *SMAD4* (∼40%)^16^, but therapeutic strategies have only been established for rare genomic alterations in <10% of patients^17^. However, recent development of KRASi and their early successes and broad applicability have garnered considerable excitement despite almost all patients developing resistance^18^. Preclinical studies suggest KRASi remodels the TME and can reinvigorate immune responses^19,20^, offering a path to synergistic combination therapies which can overcome past failure to treat stromal and immune microenvironments in PDAC^21–26^.

With cell lineages and compartments dynamically coordinating phenotypic state and therapeutic response and resistance, an integrated understanding of PDAC tumours and their complex microenvironment components at scale is necessary to inform patient stratification and precision medicine strategies. Although recent spatial transcriptomics studies of PDAC have examined sizable cohorts, most have been confined to targeted biological investigations or limited to individual or paired compartments, while protein-based spatial analyses across multiple compartments and hundreds of patients remain rare^27–30^. Overlapping molecular signatures and the fragmented nature of current studies have hindered the integration of findings into a comprehensive framework that resolves spatial co-localization and cellular heterogeneity as functional units within the PDAC TME.

To extend beyond current classification systems, we present a comprehensive spatial, genomic, and transcriptomic profile of human PDAC, providing a deeper understanding of conserved tissue contexture programs and their molecular drivers, and then linking genotype– phenotype associations to clinical outcomes. We performed high-resolution, single cell spatial analyses of 104 unique proteins in human pancreatic tumours to capture the full spectrum of TME associations and precisely quantify intra- and inter-tumoural spatial heterogeneity using imaging mass cytometry (IMC). We identified 7 epithelial subtypes that span a continuum between classical and basal phenotypes, revealing intermediate states enriched in patients with distinct clinical profiles and changes in immune, fibroblast, and endothelial cell content. Cross-compartment single cell interactions identified 8 reproducible, spatially discrete microenvironment types which neighbour and surround malignant ducts. Whole-slide laser-capture microdissection followed by mass spectrometry (LCM-MS) and scRNAseq-guided integration of proteomic data identified microenvironment- and cell-specific pathway activation. Matched whole-genome sequencing revealed associations between genomic aberrations and specific tumour subtypes, microenvironments and single cell content. Finally, using machine learning prediction models trained on our multi-omic data, we compared the predictive potential of different data types and identified non-redundant cross-omic features most predictive of overall survival, augmenting known information and highlighting key determinants of PDAC outcome.

## Results

### High-dimensional spatial profiling across PDAC tissue compartments reveals cell states associated with clinical and genomic features

To resolve the cellular architecture of PDAC, we designed and optimized three independent 40–43-plex antibody panels against epithelial, immune, and stromal cell compartments, using scRNAseq data and literature as a guide^10,31–38^. Panels were applied to consecutive serial sections of a PDAC tissue microarray (TMA) from 221 patients, ∼4 cores each (Fig. 1a; Extended Fig. 1–4; Supplementary Table 1; Supplementary Table 2). 2473 multiplexed images were acquired by IMC to identify cancer cell phenotypes, lymphoid and myeloid lineages and stromal cells, (Fig. 1a; Supplementary Table 2). Images were segmented using Deepcell^39^ into 10.1 million single cells, clustered using Phenograph^40^, and then manually annotated (Fig. 1b–c, 1e, 1g). Each compartment followed a distinct annotation strategy and biologically similar clusters were grouped together (Extended Fig. 2–4). This resulted in 7 distinct epithelial cell types, 37 immune populations, and 37 stromal populations (26 CAF/mural and 11 endothelial cell types).

**Fig 1:**
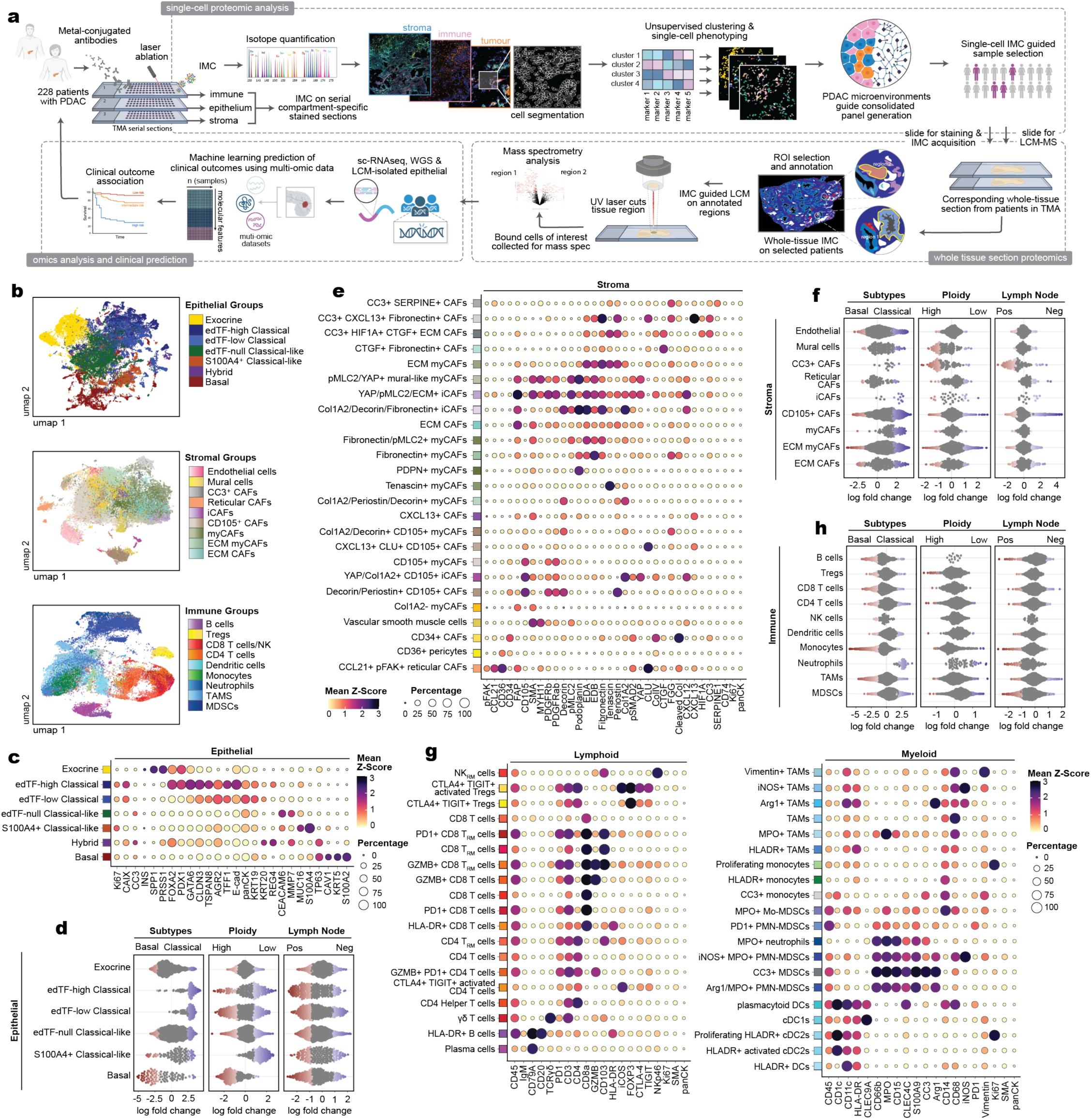
*In situ* tumour, immune and stromal single cell phenotypes are associated with genetic and clinical parameters. **a**, Schematic illustrating the multi-omic project workflow. IMC was performed on three serial sections of a TMA from 228 patients with PDAC. Five-micron IMC on whole tissue sections from corresponding patients in the TMA were stained using a consolidated cross-compartment isotope-tagged antibody panel to guide LCM on distinct microenvironments. Mass-spectrometry, bulk and scRNAseq, and whole genome sequencing were performed to further explore these microenvironments. IMC, imaging mass cytometry; TMA, tissue microarray; PDAC, pancreatic ductal adenocarcinoma; scRNAseq, single-cell RNA sequencing; CyTOF, cytometry by time of flight; LCM, laser capture microdissection; ROI, region of interest; WGS, whole genome sequencing. **b,** UMAP projection of analyzed epithelial cells (top), stromal and endothelial cells (middle), and immune cells (bottom). The legend colours correspond to general cell categories. edTF, epithelial differentiation transcription factor, CC3, cleaved caspase 3; CAF, cancer associated fibroblast; iCAF, inflammatory; myCAF, myofibroblastic; ECM, extracellular matrix; **NK,** natural killer; TAM, tumor associated macrophage; PMN-MDSC, polymorphonuclear-myeloid derived suppressor cell; DC, dendritic cell. **c,** Average expression of markers across our seven defined epithelial cell types from PDAC samples stained with the epithelial panel of isotope-conjugated antibodies. For bubble plots, the circle represents the proportion of cells that express each corresponding marker above a Z-score of 1. Markers were Z-scored and the colour represents the average Z-score for each marker per cell type with visualization capped at O and 3. **d,** Beeswarm plots of log2 fold change between binary categories **(RNA** subtype, ploidy, and lymph node status) in unsupervised neighborhoods stratified by epithelial tumour type. Differential abundance neighborhoods at FDR 10% are coloured as per Dann et al. High ploidy is >2, low ploidy is <2. FDR, false discovery rate. e, Average expression of markers across CAF and mural cell types from PDAC samples stained using the stromal panel of antibodies. CAF, cancer associated fibroblasts. **f and** h, Beeswarm plots of log2 fold change between binary categories (RNA subtype, ploidy, lymph node status and OS) in unsupervised neighborhoods stratified by stromal populations (f) and immune populations (h). Differential abundance neighborhoods at FDR 10%. g, Average expression of lineage markers across lymphoid and myeloid cell types from PDAC samples using the immune panel of isotope-conjugated antibodies.

PDAC classical and basal subtype frequencies have been estimated at about 80% and 20% in resected tumours, respectively^41^, but a substantial proportion of cells exist in a transcriptional grey zone^10,12,13,42^. Through our protein-based approach, we captured four phenotypic variations of classical cells with distinct expression and relationship to outcomes (Fig. 1c; Extended Fig. 2). The primary differentiator was tiered expression of GATA6, FOXA2 and PDX1, known pancreatic epithelial differentiation transcription factors (edTFs) linked to the classical program^6,42–44^ (Extended Fig. 2d–e). Of tumour cells 26.7% were edTF-high classical with strong expression of all three edTF, 13.6% of cells were edTF-low classical with at least one classical edTF, and 36.7% of cells were edTF-null classical-like lacking almost all basal or classical markers, but expressing non-canonical markers (CEACAM6, MMP7, MUC16), possibly representing a population of “intermediate” malignant cells as described in other papers^13,42,45^. CEACAM6, the most enriched protein in the edTF-null classical-like category, promotes EMT, is associated with poor prognosis in PDAC, and is observed in classical RNA signatures and progenitor proteomic signatures^6,42,46^. Along the continuum between phenotypes, we additionally observed an S100A4+ classical-like group (8.1%, S100A4+ without basal markers). Although S100A4 is linked to EMT and fibrosis, these Phenograph clusters lacked canonical basal markers and displayed some co-occurrence with classical marker expression in both scRNAseq and IMC data (Extended Fig. 2a, 2e), so were similarly assumed to be a ‘classical*-*like’ subtype. The remaining 15% of tumour cells were split between cells expressing canonical basal markers (11.8%, TP63/KRT5/S100A2), and hybrid cells with both basal and classical markers (3.2%, TP63 and GATA6/FOXA2/PDX1 co-expression) (Fig. 1c).

To evaluate clinical relevance of these expanded tumour cell types, we applied MILO to test for differential abundance using K-nearest neighbour (KNN) graphs of cell phenotypes across binary patient classification categories, including tumour transcriptional subtype, ploidy, and lymph node involvement^47^. Basal cells were associated with high ploidy whereas S100A4+ classical-like cells were enriched in low-ploidy tumours (Fig. 1d), reinforcing the classification of these S100A4+ cells as a unique phenotype.

In the stromal compartment, we annotated diverse CAF populations, including CD105+ CAFs, myofibroblastic CAFs (myCAFs), inflammatory CAFs (iCAFs), ECM-embedded CAFs (ECM CAFs and myCAFs), mural cells, along with diverse endothelial cell types confirming and merging independent CAF classification systems (Fig. 1e; Extended Fig. 3a,d–h). Broadly, myCAFs represented the dominant CAF population (Extended Fig. 6b), and they were enriched in tumours with a classical RNA subtype in MILO analysis, while cleaved Caspase 3+ (CC3+) cells were enriched in tumours associated with detected lymph node metastases, and CD105+ CAF populations trended towards enrichment in classical transcriptional subtype tumours (Fig. 1f).

Immune cell analysis quantified lineage-specific marker expression across a broad range of lymphoid and myeloid populations (Fig. 1g; Extended Fig. 4); the largest proportion of immune cells was composed of tumour associated macrophages (TAMs) and monocytes, with non-regulatory CD4+ T cells (hereafter referred to as CD4+ T cells) representing the dominant lymphoid subset (Fig. 1g; Extended Fig. 6a). TAMs and monocytes were associated with basal transcriptional signatures, whereas neutrophils and myeloid derived suppressor cells (MDSCs) were more linked to classical signatures (Fig. 1h). Notably, MDSCs were associated with low tumour ploidy and positive lymph node status, defining an immunosuppressive phenotype correlated to poor prognosis, independent of ploidy. Together, these data define 81 distinct cell populations in the epithelial–immune–stromal landscape of PDAC, highlighting specific cell states that are enriched in the established transcriptional subtypes and associate with genomic and clinical features, but have no clear relationships with current patient classification.

### Expanded PDAC epithelial subtypes exhibit distinct prognostic associations and immune– stromal compositions

Tumour types were assigned according to the most abundant epithelial cell annotation in each sample (Fig. 2a). Patients whose tumours were predominantly composed of exocrine cells were reassigned to their next most abundant epithelial phenotype to avoid classification based on tumour-associated normal tissue (Extended Fig. 5a–b). Most tumours were classified as classical-dominant (187/221; 84.6%), followed by basal-dominant (25/221; 11.3%) and a small subset exhibited a dominant hybrid epithelial phenotype (7/221; 3.2%). All tumours displayed mixed compositions, and distinct classical subgroups dominated by different classical cell types were apparent (Fig. 2a). Basal, hybrid, and edTF-null classical-like cells were each associated with elevated hazard as their proportion increased (Fig. 2b; Cox regression, p = 0.034, p = 0.0045, and p = 7.56e^-5^, respectively). Kaplan–Meier (KM) analysis confirmed survival stratification of patient subgroups defined by dominant epithelial phenotype with basal and S100A4+ classical-like patient groups associated with significantly worse overall survival compared to edTF-high classical dominant patients (Fig. 2b; log-rank test, p = 0.045 and 0.022 respectively). These clinical associations highlight the prognostic relevance of multiplexed imaging-defined malignant cell states and support the idea that the proteomic single cell phenotypic spectrum of tumour cells stratifies patients by differing levels of clinical risk.

**Fig 2:**
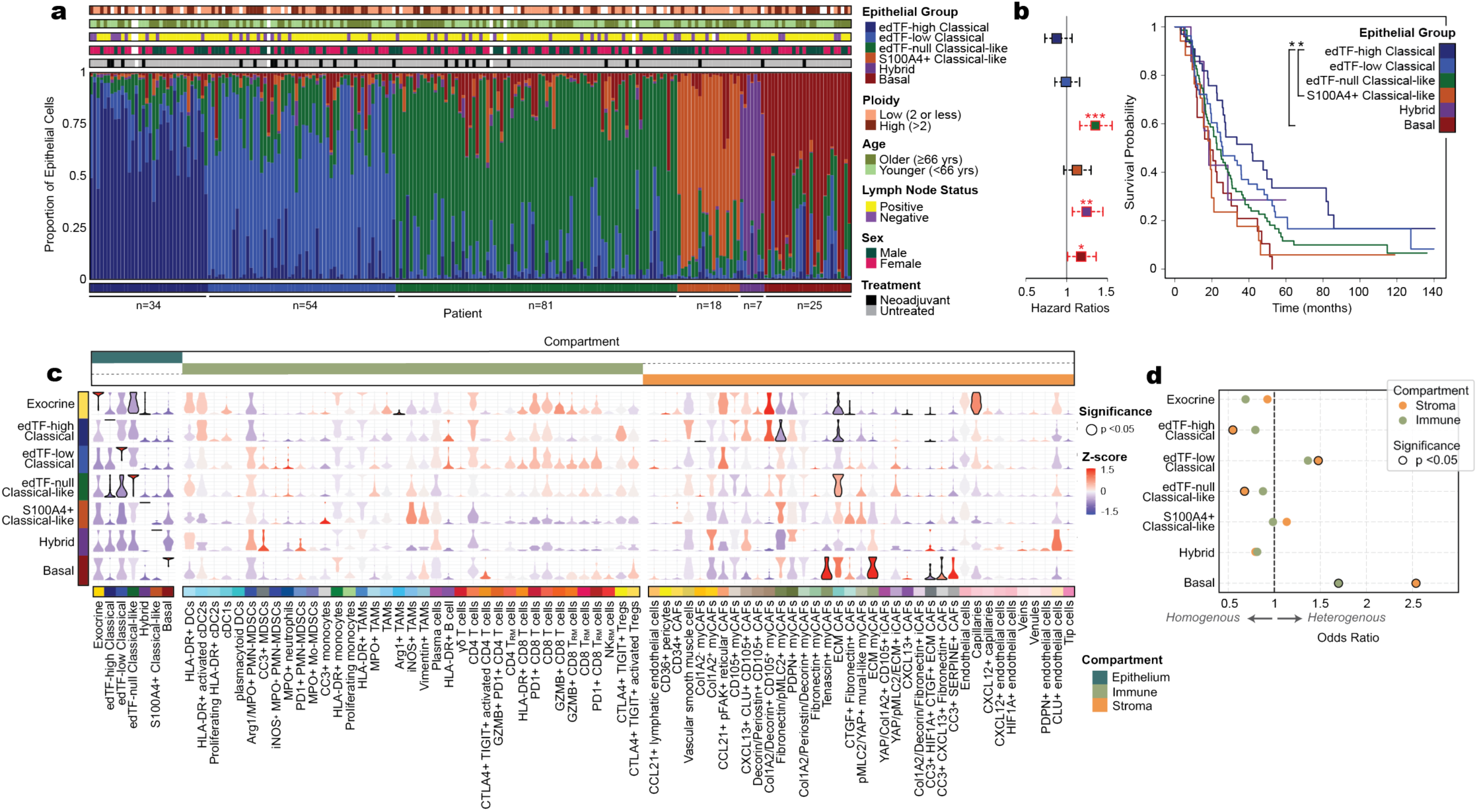
Expanded PDAC tumour subtypes with differing survival are associated with diverse immune and stromal landscapes. **a**, Stacked bar plot showing the proportional composition of epithelial cell subtypes for each tumour in which epithelial cells were captured (n=219; two patients without epithelial data excluded). Each bar represents a single patient, with the y-axis indicating the relative abundance of each epithelial population. Tracks above epithelial bars display various clinical parameters associated with each patient. **b,** Hazard ratios (left) for each *epithelial* tumour type, calculated using Cox proportional hazards regression. Points outlined in red are statistically signficant. Asterisks above each point indicate significance level: Basal: P=0.034(*), Hybrid: P=0.005(**), edTF-null Classical-like: P<0.00001(***). Kaplan-Meier OS curves (right) for each dominant *malignant* epithelial tumour type across 221 patients. Statistical significance was assessed using log-rank test highlighting significant survival differences between edTF-high Classical v. Basal P=0.022(*) and edTF-high Classical v. S100A4+ Classical-like P=0.045(*). OS, overall survival. **c,** Violin plot showing the proportion of each independent immune, stromal and epithelial cell type across patients, grouped by their dominant epithelial tumour type. The outline shade of each violin represents the statistical significance, while the colour represents deviation from the cell type’s average within each tumour type. For each cell type, we compared its proportion in each tumour type to its proportions in all other tumour types using the Mann-Whitney U test. d, Forest plot showing odds ratios from logistic regression analyses comparing the likelihood of immune or CAF heterogeneity in each epithelial subtype against all other epithelial groups. Each row represents one epithelial subtype, with immune (green) and CAF (orange) odds ratios plotted side-by-side on a shared axis. Dot size and outline intensity reflect statistical significance based on raw p-values from the logistic regression model (two-sided; outline intensity corresponds to p <0.05, 0.01, 0.001, and 0.0001). Odds ratios greater than 1 indicate higher likelihood of heterogeneity for the specified compartment in that epithelial subtype relative to others.

As such, we examined the immune and stroma composition of these epithelial subtypes, revealing distinct differences between their tumour microenvironments (Fig. 2c; Extended Fig. 6c–d). Tumour regions dominated by exocrine, edTF-high classical, and edTF-low classical phenotypes showed increased lymphocyte infiltration, namely activated and tissue resident memory CD8 T cells (CD8 TRM cells), revealing a link between elevated expression of pancreatic edTFs and increased engagement of the adaptive immune arm. This was characterized by a pronounced increase in CD4+ T cells, the most abundant lymphocyte population in our dataset, alongside antigen presenting HLA-DR+ B cells (Fig. 2c; Extended Fig. 6a). In addition to immune enrichment, endothelial cells, especially capillaries (Extended Fig. 3f; *p* < 0.001), were more frequently associated with exocrine and edTF-high classical tumours, likely reflecting increased metabolic support, altered immune cell trafficking, and greater exposure to systemic therapeutic agents. Across the classical subtypes, decreasing edTF expression was accompanied by increasing MPO+ MDSCs, neutrophils, and CAF populations with more complex matrix and phosphorylated myosin light chain 2 (pMLC2) expression, suggesting physical stiffness or tension. Hybrid tumours contained MDSCs like poor-outcome classical tumours, but even lower T cell infiltration. At the far end of the spectrum, tumours dominated by S100A4+ classical-like and canonical basal phenotypes exhibited a higher proportion of Vimentin+ TAMs, a mesenchymal-like immune population which promotes EMT, immune suppression, and matrix remodeling within the TME of solid tumours^48^. These subtypes also showed the lowest lymphocyte infiltration and significantly higher association with tenascin+ myCAFs and ECM-embedded myCAFs (Fig. 2c; Mann-Whitney test, p = 0.0067 and 0.0012, respectively).

To assess spatial cell–cell interactions, we performed permutation-based pairwise analyses, revealing compartment-specific preferences for homotypic associations—epithelial with epithelial, immune with immune, and CAF with CAF—consistent with patterns reported across cancer types^49,50^ (Extended Fig. 7b). Within-compartment interactions reflected the immune cell content of each subtype; for example, classical tumours with any edTF expression showed significantly more CD8 and CD4 T cell interactions than would be expected by random chance, deemed “attraction”, coinciding with relatively increased immune infiltration (Fig. 2c; Extended Fig. 7a). Across the entire dataset, distinct immune cell interaction patterns emerged: B cells, CD4 T cells, CD8 T cells, and DCs were mutually attracted. CD4 T cells exhibited specific affinity for Tregs, while DCs were preferentially attracted to TAMs, suggesting potential cell-specific immunosuppression. Additionally, neutrophils and MDSCs were attracted (Extended Fig. 7a). Within the CAF compartment, CD105+ CAF were spatially proximal to immune and endothelial cells, while ECM myCAFs had significantly fewer interactions with these populations than random chance, aligning with observed infiltration trends (Fig. 2c; Extended Fig. 7d). ECM CAFs, ECM myCAFs and CC3+ CAFs lacked interactions with each other and other cell types, especially in basal tumors where they were highly enriched (Extended Fig. 7b, 7d).

Despite the clear differences in cellular content and interactions between tumour types, considerable intra-patient regional heterogeneity was observed, with cored samples from different locations enriched for distinct cell types and spatial arrangements (Extended Fig. 6c–d). Thus, we used Kullback-Leibler (KL) divergence to quantify differences in the distribution of cell content between the ∼4 imaged regions from every patient (Extended Fig. 6e). In epithelial, immune, and stromal cell compartments approximately half of the tumours were spatially homogenous and < ∼10% of patients had extensive spatial phenotypic heterogeneity, with the most observed in immune and stromal compartments (Extended Fig. 6e). When comparing tumour types, edTF-high classical tumours exhibited a significantly more uniform stromal composition (Wald test, p = 0.041) and a trend towards more homogenous immune composition, whereas edTF-low classical tumours showed significantly higher heterogeneity in both compartments (Fig. 2d; Wald test, stroma: p = 0.0473; immune: p = 0.005). Basal-dominant tumours displayed the greatest stromal and immune heterogeneity (Fig. 2d; Wald test, both p < 0.0001).

Overall, our data refine the classical–basal continuum and link it to TME content in primary tumours, ranging from lymphocyte-rich, vascularized milieus to progressively immunosuppressive, ECM-dense stroma as classical programs wane and basal programs predominate, with the latter also exhibiting greater inter-regional heterogeneity.

### Recurrent cross-compartment cellular microenvironments are associated with patient survival

To more comprehensively characterize how epithelial, stromal, and immune elements co-organize locally as multi-cellular PDAC microenvironments, we performed spatial alignment and cross-compartment analyses independent of molecular subtype by using epithelial morphology as a fiducial on which to register our three serially stained TMA sections (Fig. 3a, Methods). All cells were assigned to their nearest epithelial ‘lesion’ matched across sections (Fig. 3a; Supplemental Table 3) and clustered based on cell type composition, enabling reconstruction of eight reproducible, spatially distinct, multi-cellular tumour microenvironments that integrate all three compartments (Fig. 3b, left heatmap; Extended Fig. 8a). Microenvironments included: 1. *Immune suppressed:* driven by immunosuppressive myeloid lineages such as TAMs and polymorphonuclear myeloid-derived suppressor cells (PMN-MDSCs); 2. *ECM-rich:* fibroblast populations embedded in complex ECM including tenascin and fibronectin; 3. *Stiff matrix:* stroma with pMLC2 (Extended Fig. 8c) resulting from matrix tension and biophysical force; *4. Immune-infiltrated stroma:* myeloid cells and lymphocytes within myCAF and iCAF-rich regions; 5. *CD105⁺ CAF high*: enriched for collagen- and decorin-embedded CD105⁺ myCAFs; 6. *CD105⁺ fibrovascularized*: highly vascularized microenvironments dominated by CD105⁺ CAFs and infiltrated by CD4⁺ and CD8⁺ T cells; 7. *Immune infiltrated*: densely populated by 33 immune cell types from both myeloid and lymphoid lineages; and 8. *Collagen-rich myCAF*: dominated by collagen-expressing myCAFs (Fig. 3b, 3d; Supplemental Table 3).

**Fig 3:**
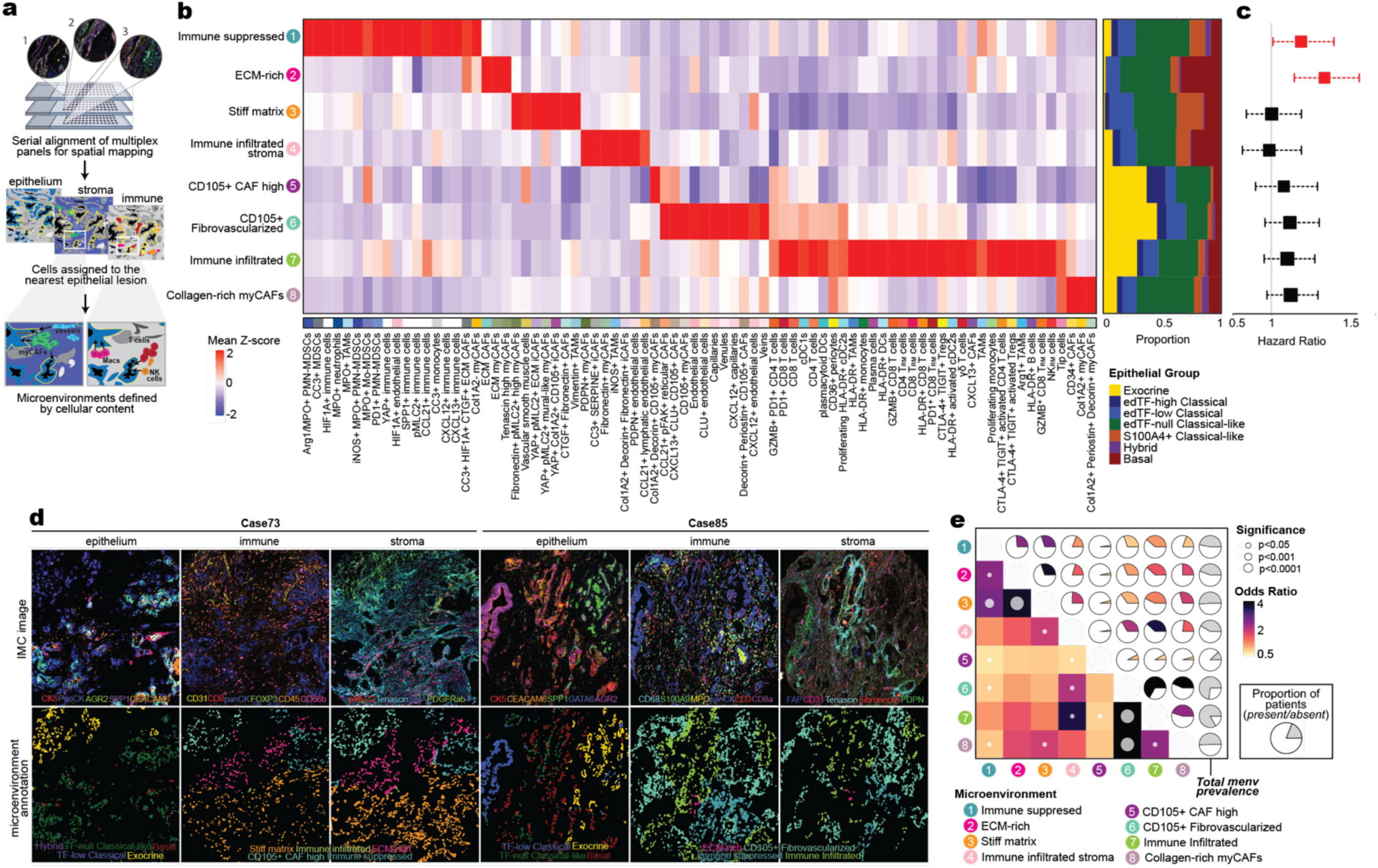
Reproducible spatial relationships define outcome associated tumour microenvironments. **a**, Schematic illustrating the computational alignment of serial TMA sections stained with custom antibody panels targeting epithelial (section 1), stromal (section 2) and immune (section 3) compartments. TMA, tissue microarray. **b**, Heatmap showing representative cell population expression across microenviroments (left) and the crorresponding proportion of neighbouring epithelial lesions for each microenvironment (right). **c**, Hazard ratios for each microenvironment where eachpatient was assigned dominant microenvironment; hazard ratios were derived using Cox proportional hazards regression. Red outlines indicate statistical significance (p < 0.05). **d**, Representative multiplexed IMC images from Case 73 and Case 85 across epithelial, immune, and stromal compartments (top), and corresponding microenvironment annoations generated using cytomapper (Eling et al., Bioinformatics, 2021) on aligned serial sections (bottom).**e**, Co-occurence and prevalence of microenvironments across patients. Upper triangle pie charts show co-occurence frequency between eachpair of microenvironments. Wedge size denotes patients with both microenvironments as a proportion of the entire cohort (n=221). Colour denotes odds ratio. Lower triangle, ties show odds ratios with significance (p<0.05) indicated by white circles scaled by p-value. Rightmost column indicates overall prevalence per microenvironment.

Identifying the epithelial subtype composition within each microenvironment revealed preferential spatial relationships. Basal epithelial lesions, morphologically identifiable as poorly differentiated clusters of epithelial cells, were commonly interwoven within *ECM-rich* stromal microenvironments (Fig. 3b, right), corresponding with previous work^51^. In contrast, both *CD105*+ *CAF high* and *CD105*+ *fibrovascular* microenvironments were frequently adjacent to exocrine and edTF-high classical lesions (Fig. 3b, right). Further analysis revealed that 12/13 patients with predominant *CD105*+ *high CAF* microenvironment had been neoadjuvantly treated, even though only 26/221 patients in our cohort received neoadjuvant therapy, and it was much more enriched in platinum-treated tumours than in radiation- or non-platinum chemotherapy-treated (Extended Fig. 8b; Supplemental Table 1). Specifically, it was the dominant microenvironment in 8/9 patients treated with platinum-based chemotherapy, comprising 86.46% of all microenvironments regardless of epithelial subtype and expressing significantly elevated fibrinogen (FGG; Fig. 3b; Extended Fig. 8d–f), a key glycoprotein involved in coagulation, wound healing, ECM remodeling, and inflammation. These findings demonstrate that, compared to untreated TMEs, platinum-induced tissue remodeling promotes a pro-fibrotic stromal niche defined by Col1A2/Decorin high CD105+ CAFs with implications for therapeutic response and microenvironmental reprogramming in the post-treatment setting.

Immune cells, when present within basal-dominant tumours, were primarily localized to *immune-infiltrated stromal* microenvironments (Fig. 3b), enriched for two populations known to induce cytotoxic inflammation: iNOS+ TAMs, an M1-like macrophage population, and HLA-DR+ activated DCs, which present antigens to CD4+ T cells. These likely act in concert with surrounding iCAFs to mediate localized anti-tumour effects and were significantly associated with *immune-infiltrated* microenvironments (Fig. 3e; odds ratio = 3.51). In contrast, *ECM-rich* stromal microenvironments devoid of immune infiltration, often associated with basal-dominant tumours, were linked to significantly worse clinical outcomes (Fig. 3c; hazard ratio (HR) = 1.34, 95% confidence interval (CI): 1.15–1.55; Cox proportional hazards model, p = 0.00015). Similarly, *immune suppressed* microenvironments, characterized by the presence of granulocytic or PMN-MDSCs and TAMs, were associated with significantly increased hazard (Fig. 3c; HR = 1.19, 95% CI: 1.01–1.39; Cox proportional hazards model, p = 0.035), and often co-occurred with *ECM-rich* and *stiff matrix* microenvironments, suggesting spatial or functional coupling between these compartments (Fig. 3e; odds ratio = 2.46 and 2.64, respectively). Protein marker expression analysis revealed the highest hypoxia-inducible factor 1-alpha (HIF1A) expression in *immune suppressed* microenvironments relative to all other microenvironments, along with elevated expression of neutrophil extracellular trap (NET) affiliated myeloperoxidase (MPO; Extended Fig. 8c). This supports the notion that beyond their known immunosuppressive roles in PDAC, PMN-MDSCs in hypoxic, MPO-rich niches, may also contribute to oxidative tissue damage, fostering a tumour-permissive environment that promotes progression.

Collectively, this represents an overarching framework of TME relationships in PDAC associated with variations in outcome. Exocrine, edTF-high and edTF-low classical lesions were more commonly associated with *immune-infiltrated* and *fibrovascularized* niches defined by CD105+ CAFs and increased vasculature. In contrast, basal tumours were frequently surrounding *ECM-rich* and *stiff matrix* environments enriched in iCAFs and pMLC2, alongside *immune suppressed* innate microenvironments with reduced lymphocytes in the same patient.

### Integrated spatial proteomics and single cell transcriptomics identify environment-specific cellular functions

To expand our understanding of these tumour subtypes and eight microenvironments, we consolidated our epithelial, immune, and stromal panels into a single 42-marker IMC panel designed to provide us with sufficient information to identify these unique regions and performed whole-slide multiplexed imaging at 5 µm resolution (Fig. 4a; Extended Fig. 9; Supplementary Tables 4, Methods). In select patients representing all PDAC disease states identified in our TMA, we visualized patient-level spatial heterogeneity in full tumour sections, confirming the associations identified in sub-sampled TMAs (Extended Fig. 9; Supplementary Table 5). Even within a single heterogeneous tumour, tumour–immune–stroma relationships varied across spatial scales, with immune hot and cold niches embedded in a broader mixed stroma neighbouring classical or basal subtype lesions. To complement our original single cell IMC profiling, mass spectrometry (MS)-based proteomics achieved deeper, unbiased molecular readouts of each spatial niche, capturing ECM components and signaling networks that shape these environments. Guided by multiplexed imaging, between 12–33 representative regions per tumour subtype and microenvironment all from >3 patients were collected by laser capture microdissection (LCM) followed by MS (n = 11 patients, n = 275 regions/samples; Fig. 4b, left heatmap; Supplementary Table 6; Methods). An average of ∼3500 proteins were identified per epithelial lesion or microenvironment and these were integrated with single cell RNA expression data from a collection of eight publicly available datasets (n = 163; Fig. 4b, right heatmap; Supplementary Table 6).

**Fig 4:**
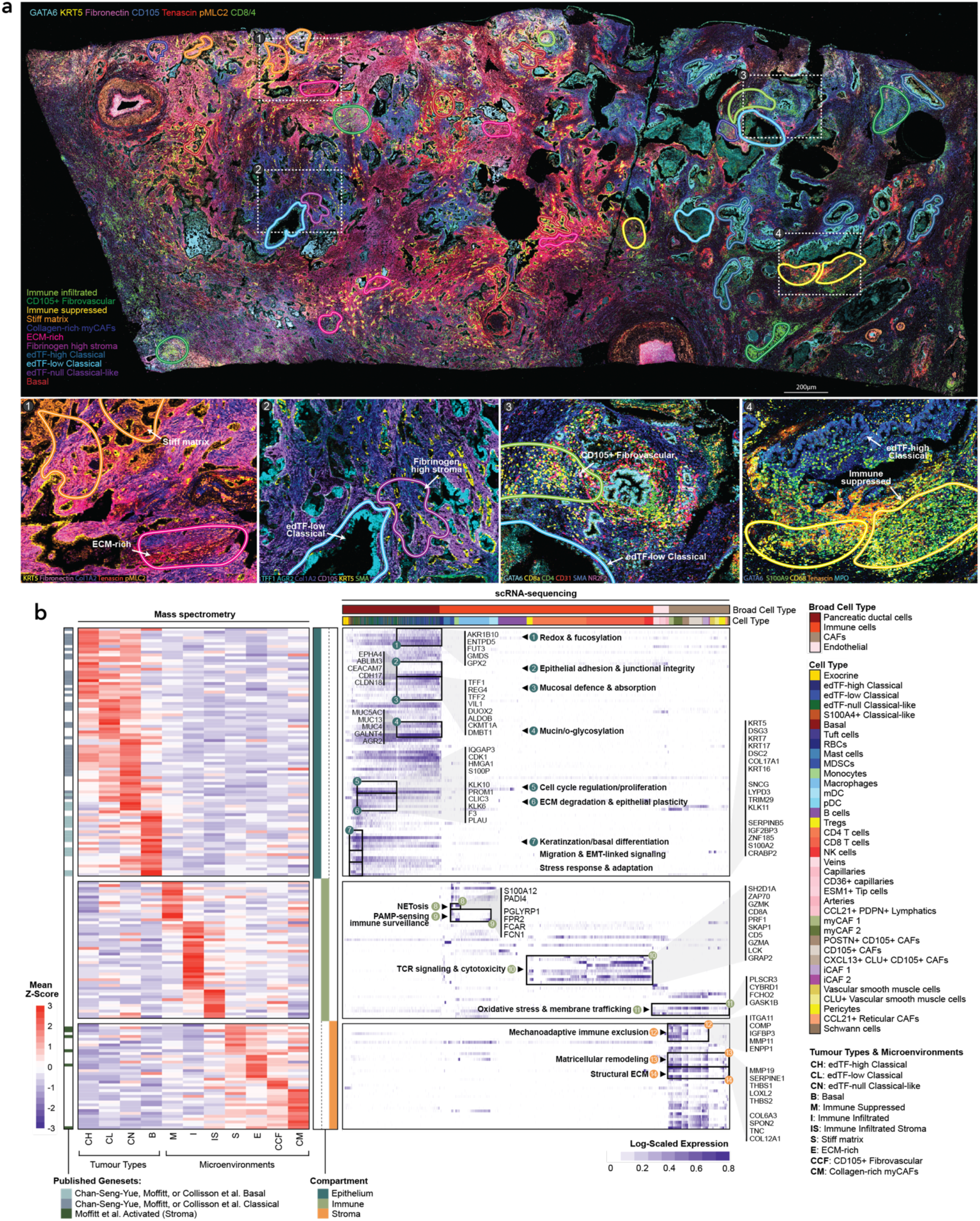
Spatially resolved epithelial sybtypes and microenvironments exhibit functional signatures across multi-omic data. **a**, Whole-issue PDAC section imaged by IMC at a 5µm resolution showing spatially defined microenvironments characterized by epithelial, immune, and stromal markers from our consolidated cross compartment panel. Annotated microenvironments represent combinations of distinct epithelial tumour types and representative microenvironments. Insets highlight representative regions with dominant features such as 1) ECM-high CAFs and matrix stiffness (1), fibrinogen high stromal zones, (2), CD105+ fibrovascularized (3), and locations of immune-suppression (4). The full resolution images will appear in the final publicatio. IMC, imaging mass cytometry; ECM, extracellular matrix. **b**, *Left*: Heatmap of average protein abundance across laser capture microdissection (LCM) regions profiled by mass spectrometry (LCM-MS), corresponding to the microenvironments annoated in (a). Columns represent individual microenvironments; rows represent proteins, z-score normalized across all regions. *Right*: Heatmap of expression of genes encoding the proteins identified by LCM-MS, as detected in single-cell RNA-seq data and stratified by IMC-defined microenvironments. Columns represent individual cells, annoted using SingleR based on known marker genes. Gene expression is log-normalized and row-scaled to the maximum expression per gene. ECM, extracellular matrix; CAFs, cancer-associated fibroblasts.

Cross-validation of markers that overlapped across multiplexed imaging, proteomic, and scRNAseq datasets confirmed accurate detection of target cell populations and identification of mast cells not measured by IMC, specifically in *immune infiltrated* microenvironments (Extended Fig. 10a–b). edTF-high classical tumours retained junctional integrity and ductal organization marked by elevated CDH17, CLDN18, and CEACAM7. Concomitant upregulation of REG4 and DMBT1, implicated in goblet/enteroendocrine precursor maintenance and barrier immunity at the tumour front, is consistent with the description of classical PDAC as a gastrointestinal progenitor-like state^9,52^. This is further supported by AKR1B10 which facilitates lipid metabolism and detoxification and the antioxidant GPX2, both mitigating ROS-specific oxidative stress and supporting high metabolic and secretory demands of well-differentiated classical tumours^53,54^. Similar to previous findings in classical tumours, edTF-high classical tumours also showed increased expression of fucosylation enzymes (FUT3 and GMDS), enhancing fucosylated glycans on mucins, adhesion molecules, and receptors, thereby altering cell–cell and cell–ECM binding, immune interactions, and receptor signaling^55,56^. But, as tumour cells shift from edTF-high to edTF-low classical subtypes, a decrease in edTFs is accompanied by a shift toward a secretory, mucin-rich phenotype. edTF-low tumours contain considerable MUC4, MUC13 and MUC5AC, heavily glycosylated proteins, and GALNT4, a critical initiator of mucin-type O-glycosylation, that form a protective mucus barrier known to shield tumours from immune surveillance, compromise adherence and polarity, facilitate metastatic dissemination, and alter receptor-mediated signaling^57^. In particular, MUC5AC, typically restricted to gastric epithelium, is a well-documented component of the mucus hypersecretion program observed in PDAC^58^. These represent distinct yet related glycosylation programs along the epithelial spectrum.

Within our cohort of resected PDACs, conventionally known to be immune-cold, edTF-high classical tumours contain the most T cell enriched *immune infiltrated* microenvironments, with increased TCR signaling (elevated ZAP70, LCK, GRAP2, SKAP1, SH2D1A) and cytotoxic effectors (PRF1, GZMA, GZMK), suggesting T cells are not just present but functionally engaged (Fig. 4b)^59^. These microenvironments contained limited myeloid activation but often co-occurred with *immune infiltrated stroma* microenvironments (Fig. 3e) where macrophages displayed increased *FCHO2*, enabling clathrin-mediated endocytosis (CME), highlighting an active scavenging and antigen-processing role (Fig. 4b)^60^. This environment also contained CYBRD1, a regulator of iron handling and ferroptosis sensitivity, and PLSCR3, involved in lipid remodeling and apoptotic signaling, indicating redox and membrane dynamics optimized for antigen uptake and metabolic adaptation^61,62^. Dampened immune infiltration, increased immunosuppression, and altered ECM and CAF content was associated with decreasing edTF expression across the classical subtypes (Fig. 2c), and proteomics further revealed the localized activity and function of these environments.

At the aggressive end of the classical spectrum, edTF-null classical-like tumours displayed coordinated programs driving ECM degradation, invasion, proliferative expansion, and metastatic potential. This included ECM remodeling proteins CLIC3 and PLAU, as well as KLK6 and KLK10, serine proteases that activate growth signaling, degrade ECM, and facilitate EMT and metastatic spread (Fig. 4b)^63,64^. This is accompanied by heightened cell cycle activity marked by G2/M cell cycle checkpoint CDK1 and scaffolding protein IQGAP3 (G1/S), along with increased cancer stem cell marker PROM1, suggesting stem-like functions known to be associated with metastasis and therapy resistance in PDAC^65^. Interestingly, high expression of HMGA1, a chromatin remodeling protein that promotes therapy resistance and dedifferentiation to a stem-like state was also detected^66^. These programs jointly define poorly differentiated, ECM degrading and invasive edTF-null classical tumours.

At the far end of the microenvironment axis, NETosis and innate immune myeloid activity are localized to *immune suppressed* microenvironments and hypoxic regions. This includes neutrophil and PMN-MDSC-derived factors, including *PADI4,* required for chromatin decondensation during NET formation, and the pro-inflammatory alarmin *S100A12*^67^ (Fig. 4b), along with MPO marker expression observed by IMC (Extended Fig. 8c). TAMs in *immune suppressed* microenvironments, unlike TAMs found in *immune infiltrated stroma*, displayed pathogen-associated molecular pattern (PAMP) sensing (*PGLYRP1, FPR1, FCN1*), suggesting a shift away from classical antigen presentation toward chronic inflammation and immune evasion^68,69^. Our data demonstrates myeloid cell abundance was broadly observed in tumours as expected, but spatial data revealed unique context-dependent functional states, where TAMs vary according to their local microenvironment. EdTF-null tumours had more *immune suppressed* environments and these were more likely to co-exist with *stiff matrix* and *ECM-rich* environments (Fig. 2c, 3e).

Across the subtypes, from high to low edTF expression in classical tumours and into S100A4+ classical-like, hybrid, and basal tumours, immune responses were dampened, and ECM and CAF content was altered (Fig. 2c). Basal PDAC tumours exhibited the strongest relationship with *ECM-rich* and *stiff matrix* environments which excluded many immune cell subsets. These tumours were characterized by enrichment of basal/myoepithelial cytokeratins (*KRT5, KRT7, KRT16, KRT17*). IGF2BP3, an RNA-binding protein enriched in basal tumours, stabilizes oncogenic *KRAS* and *MYC* transcripts and has been associated with transcriptional programs favouring immunosuppressive cytokines and checkpoint expressionwan^70–72^, tying basal identity to oncogene-driven immune evasion. These tumours had increased TRIM29, which promotes STING degradation, primarily limiting type I interferon (IFNI) response, as well as type III interferon (IFN-λ), dampening cytokine and chemokine production critical for cytotoxic T cell recruitment, while also reducing MHC class I expression^73^, providing a potential mechanism for the reduced T cell infiltration observed in basal tumours in both our data and a recent large-scale scRNAseq atlas^45^. Further, basal tumours expressed the retinoic acid-binding protein CRABP2 and retinoic acid dehydrogenase DHRS9^74,75^, implicating retinoic acid signaling known to be associated with mechanosensory regulation and stromal modulation. *ECM-rich* regions are marked by upregulation of thrombospondin (*THBS1* and *THBS2*), which drive collagen cross-linking and activate latent TGF-β to induce ECM organization^76^. Concurrent increases in LOXL2^77^, and structural ECM components (COL6A3, TNC, COL12A1) supports matrix cross-linkage and deposition, sustaining TGF-β activation and mechanical exclusion of immune cells. This aligns with features of co-occurring *stiff matrix* which displayed increased ITGA11, COMP and IGFBP3, triggering integrin-mediated mechanotransduction and contractility that can release and amplify TGF-β/SMAD signaling. IMC data of *stiff matrix* regions validates this, expressing the highest pSMAD2 relative to other microenvironments (Fig. 4b; Extended Fig. 8c). This coincides with enrichment of ENPP1 which also degrades cGAMP^78^, further impairing STING-dependent IFNI responses and reinforcing immune suppression in regions more associated with basal tumours and other poor outcome epithelial subtypes. These findings describe the existence of a spatially coordinated tumour–immune–stroma axis, anchored in basal differentiation, with associated ECM diversity and stiffening, TGF-β signaling, and myeloid-mediated immune suppression. This program not only contextualizes the biology driving poor prognosis, but also reveals druggable targets, such as ENPP1, that may be disrupted in patients with this pathogenic immune exclusion axis.

The co-evolution of defined tumour subtypes and their neighboring microenvironments establishes a spatial framework that links the epithelial phenotypic spectrum not only to surrounding microenvironments and clinical outcomes, but also to unique functional behaviours.

### Patient matched DNA sequencing identifies genomic relationships to tumour single cell phenotypes and microenvironments

To build on the observed epithelial phenotypes and co-evolving microenvironments, we next asked whether these states are shaped by specific genomic alterations. Within 192 patients in our cohort (Extended Fig 11; Supplementary Table 7), LCM tumour-enriched genomes revealed that driver mutations were largely homogenous as expected, and most mutations other than the non-mutually exclusive ubiquitous big four (*KRAS, TP53, CDKN2A, SMAD4*) were found at a frequency of <15% (Extended Fig. 11a). However, even within the extensive genomic instability found in PDAC^79–81^, genomic aberrations revealed a similar dichotomy along tumour subtype and microenvironment phenotypic spectrums (Fig. 5a–b).

**Fig 5:**
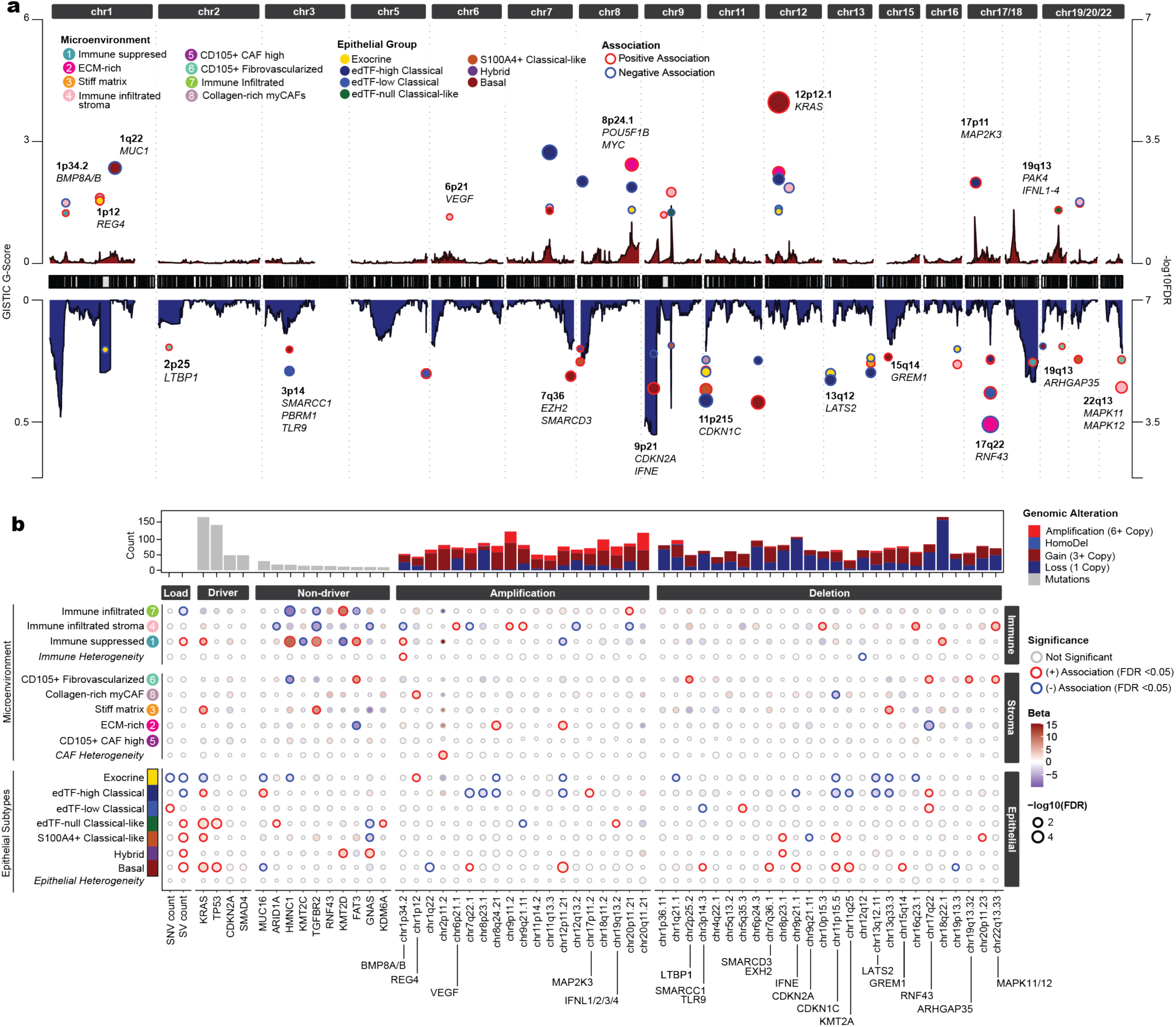
Association of whole genomes with spatially resolved tumour subtypes and microenvironments. **a**, Genome-wide associations of recurrent somatic CNAs identified by GISTIC2.0. General linear modelling was applied to test for associations between recurrent alterations and microenvironments and cell types. The results were mapped back to the corresponding cytogenetic band. Positive associations are highlighted in red and negative assoctiations in blue. Left axis represents the GISTIC G-Score, shown as red and blue density maps across the genome; Right axis represents the -log10(FDR) from the generalized linear model, mapped to genomic positions. CNA, copy number alteration. b, (Top) Bar plot representing mutational and alteration frequency of SNPs and CNAs across a curated set of features. (Bottom) Bubble plot representing associations of microenviroments and epithelial groups from a generalized linear model catergozied into genome-wide marker burden, driver and non-driver mutations and associated amplification and deletion regions called utilizing GISTIC2.0, summarized to their respective cytogenetic bands. Statistically significant associations are highlighted in red and blue (FDR < 0.05). SNP, single nucleotide polymorphism. Beta, coefficient of estimated change from linear model in logit units.

Genotype to phenotype relationships reflected the polarization of the PDAC phenotypic spectrum and its relationship to oncogenic dosage, as well as pathways associated with specific tumour types and environments. Total single nucleotide variant (SNVs) and structural variant (SVs) counts and multiple specific aberrations were negatively associated with the presence of exocrine tissue in the tumour and increased in poor outcome tumours. This included increased SNV counts in edTF-low classical tumours, SVs in all poor outcome tumours, and *KRAS* mutations broadly distributed across all tumour subtypes (Fig. 5b), however, the distribution of *KRAS* hotspot variants differed by subtype: edTF-null classical-like tumours showed a depletion of wild-type *KRAS*, whereas basal tumours were enriched for non-canonical *KRAS* variants (other than G12D, G12R, G12V or Q61H) (Extended Fig. 12g). Differential association of oncogenic drivers was evident by *12p12* (*KRAS)* amplifications and *9p21 (CDKN2A)* deletions in basal tumours, *TP53* mutations in edTF-null and basal tumours, and *11p15 (CDKN1C)* deletions in both basal and S100A4+ classical-like tumours, highlighting copy-number oncogene dosage and compounding hallmark tumour suppressor deletion as a feature of more aggressive phenotypes. These observations pair with our proteomic data revealing IGF2BP3 expression in basal tumours, which stabilize oncogenic *KRAS* and *MYC* transcripts (Fig. 4b), demonstrating that these tumours leverage multiple molecular strategies to induce hyperactive oncogenic signaling.

Genomic modifications of epigenetic regulators, genes frequently altered in a tissue-specific manner across cancer types^82^, were associated with aggressive phenotypes: *7q36* (*EZH2, SMARCD3)* and *3p14 (SMARCC1, PBRM1)* were depleted in basal tumours, *11q25 (KMT2A)* was depleted in S100A4+ classical-like tumours, *KMT2D* was mutated in hybrid tumours, and *ARID1A* and *KDM6A* loss of function mutations were associated with edTF-null classical-like tumours. Further, *13q12 (LATS2)* deletion, a regulator of both Hippo signaling and PRC2-mediated histone methylation, was negatively associated with edTF-high classical tumours. While alterations in epigenetic modifiers were primarily associated with tumour phenotype shifts, there also appeared to be an idiosyncratic association between *KMT2C/D* mutations, increased *immune infiltrated* microenvironments, and decreased *immunosuppressed* microenvironments, aligning with recent colorectal- and pan-cancer analyses^83,84^.

Beyond broad tumour spectrum shifts, we observed evidence of phenotype-specific survival mechanisms concordant with IMC and MS data. Notably, loss of *17q22* (*RNF43)*, a negative regulator of Wnt signaling, was the only genomic alteration significantly enriched in both edTF-high and edTF-low classical tumours. This deletion is expected to enhance proliferative signaling, consistent with the elevated Ki67 expression observed in these phenotypes (Fig. 1c, Extended Fig. 2e). In contrast, deletions of *9p21* (*IFNE)* and *3p14* (*TLR9)*, which impair interferon signaling^85^, were enriched in basal tumours, suggesting an alternative mechanism of suppressing immune responses that parallels the upregulation of *TRIM29* observed in basal cells by LC-MS (Fig. 4b). Likewise, *1q22 (MUC1)* amplifications were negatively associated with basal tumour phenotypes, consistent with their canonical position at the opposite end of the spectrum from the gastric, mucin-rich subtypes, which are associated with *MUC16* mutations. Together, these observations suggest that distinct genomic alterations may help prime or reinforce specific tumour phenotypes, reflecting differences in developmental context and selective pressures.

Finally, we identified genomic alterations with microenvironment-specific associations: alterations linked to immune composition were distinct from those associated with tumour phenotype, whereas stromal-associated alterations appeared to track along the tumour spectrum. Mutations in *HMCN1*, *TGFRB2*, and *FAT3* were all associated with increased *immune suppressed* microenvironments and decreased *immune infiltrated* environments (Fig. 5b). *TGFRB2* mutation was additionally associated with increased *stiff matrix*. Poor outcome *ECM-rich* microenvironments were negatively associated with *17q22* (*RNF43)* deletions, opposite to edTF-high classical tumours, and harboured *8p24.1* (*MYC)* amplifications and *12p12.1* (*KRAS)* amplifications, similar to basal tumours. This aligns with recent work showing that tenascin-high myCAFs, a hallmark of our *ECM-rich* stroma environment, are depleted in response to *KRAS* inhibitors, supporting the idea that the stromal ECM-richness has a functional dependency on *KRAS* signaling intensity^86^ (Fig 5a; Extended Fig. 12a–c). Furthermore, while links between ECM deposition, *KRAS* amplification, and basal phenotype have been observed before^10,51^, it remains an open question whether they are independent markers of progression or overlapping pathways. In our data, *KRAS* copy gain (>diploidy, n = 45), *ECM-rich* microenvironment presence (n = 38), and basal cell enrichment (>20%, n = 28), significantly overlapped between each set (p < 0.05), and across all three features (Extended Fig. 12a–c; n = 8, p < 0.0001).

Our genomic and IMC spatial analysis provided us with critical biological context on the PDAC phenotypic spectrum across epithelial states and on their surrounding TMEs, forming the foundation for leveraging and interpreting these features in a prognostic prediction model.

### Multi-omic integration using general machine learning for prognostic prediction and minimal feature selection

With 182 patients having matched genomic, clinical, and spatial single cell phenotyping, there was a unique opportunity to search for a minimal set of biomarkers defining patient outcome. To overcome the dual challenges of features > samples and inherent differences in noise between modalities, we used Stabl, a general machine-learning method that identifies robust features for multi-modal predictive models by injecting modality-specific artificial noise to create data-driven signal-to-noise thresholds for each omics dataset (Fig. 6a)^87^. Comparing prediction of survival >2 years with models derived from IMC, genomics, or clinical metadata independently, models using spatial single cell data from IMC outperformed those with only genomic features, but both provided improvements over clinical data which relied heavily on lymph node status, a variable only available after surgical intervention (Fig. 6b–c). But predictions from combined multi-modal models were best (AUROC = 0.772, Precision = 0.791) with lymph node status still providing a small but notable improvement (AUROC = 0.752 to 0.772).

**Fig 6:**
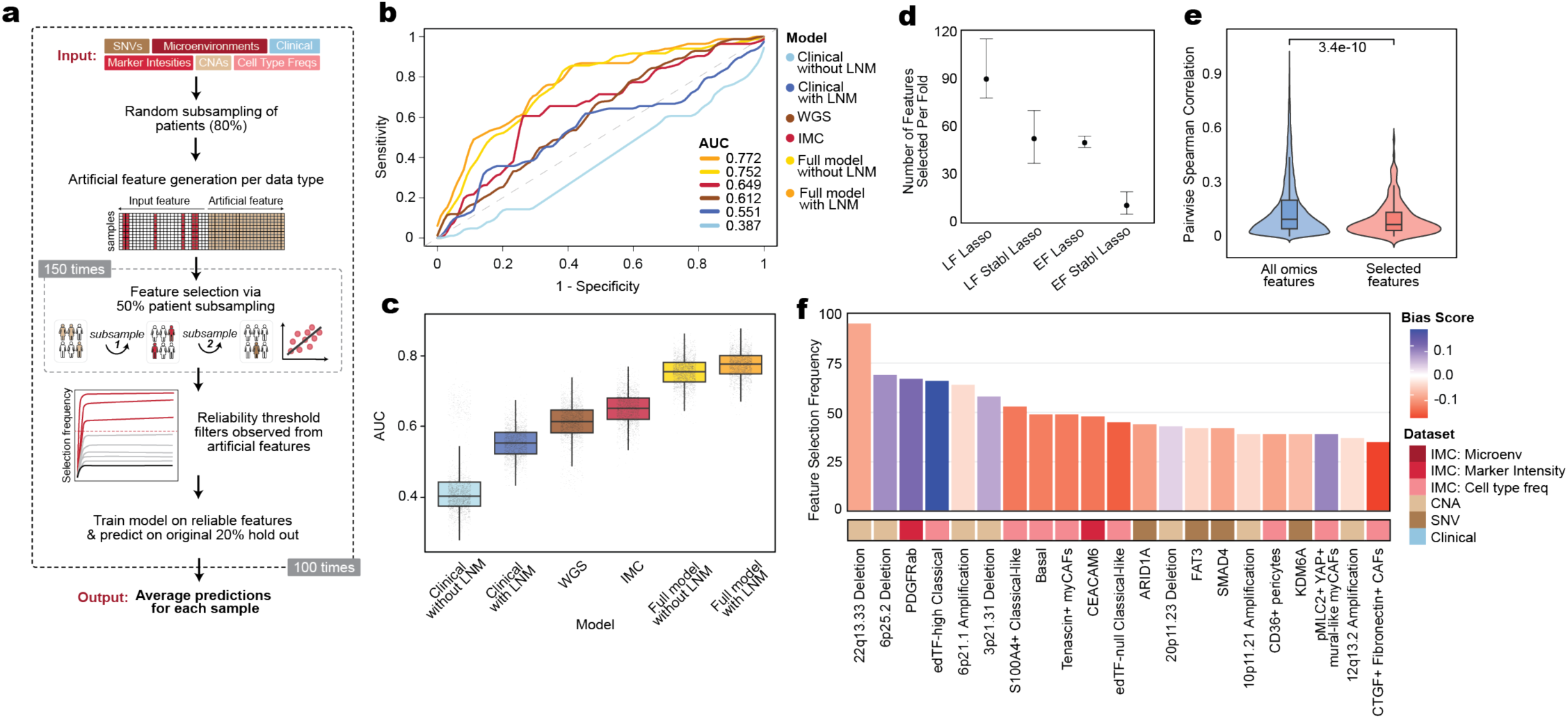
Within -omic data cell phenotype features predict overall survival. **a**, Schematic of the predictive modeling framework. Patients with matched multi-modal input features, including CNAs, SNVs, spatially normalized cell-type frequencies, protein marker intensities, and clinical variables, undergo random subsampling (80%) and then artificial feature generation. Feature selection is performed through 50% bootstrap subsampling and feature selection (n=150), with a meaningful selection frequency threshold determined by comparison to a null distribution derived from the artificial features. Features exceeding said reliability threshold are then used to train a model and predict outcomes on held-out samples (20%). The entire procedure is repeated 100 times to generate averaged predictions per patient. **CNA,** copy number alterations; SNV, single nucleotide variant. **b,** Average cross-validated AUC for models trained separately on each dataset. AUC, area under curve. c, distribution of AUC per bootstrapped model tested on cross-validation trained for each dataset. LNM, lymph node metastasis; WGS, whole genome sequencing; IMC, imaging mass cytometry. **d,** Average number of features, with 5% and 95% quantiles shown, selected by each model type across bootstraps. e, Distribution of Spearman correlation scores between all omics features (n = 82082 pairs) vs pairwise scores between EF Stabl Lasso selected features (n = 420 pairs). EF, early fusion; LF, late fusion. **f,** The top selected features across Stabl bootstrap iterations during “double fusion”, representing the most robust and informative features. The bias score represents whether a feature is associated with greater than two year survival (positive) or less than two year survival (negative).

However, each model on average utilized 92 features inhibiting interpretability and applicability. To filter to robust, non-redundant features from any source, we carried forward the multi-omic features that were selected in at least 50% of each independent technology-specific Lasso model. We then applied Stabl Lasso, which compressed the average number of features in a model to only 10.5, with an AUROC of 0.648 (Fig. 6d). The final selected features showed lower inter-feature correlation (Fig. 6e; Wilcoxon rank-sum, p < 0.0001), indicating decreased redundancy, and reflected previously observed constellations of tumour phenotypes despite reduced overall feature correlation. Basal, S100A4+ classical-like, and tenascin-high myCAFs, a key component of the *ECM-rich* microenvironment, all emerged as strong negative prognostic indicators (Fig. 6f) and were frequently linked in our spatial analyses (Fig. 2c, Fig. 3b). The second grouping of features represented a poor prognosis quartet: CEACAM6, edTF-null classical tumour phenotype, and two mutations to epigenetic regulators (*KDM6A* and *ARID1A*) that were associated with this epithelial subtype (Fig. 1c, Fig. 5b). The final pair of related features, edTF-high classical cancer cells and PDGFRɑβ expression, were associated with favourable outcomes. PDGFRɑβ was highest in the *CD105+ CAF high* microenvironment that had been observed to be enriched around edTF-high classical lesions (Fig. 3b, Extended Fig. 8d, right). Immune features were not retained as prognostic predictors. While unexpected, given the established prognostic role of immune infiltration in many cancers, immune content may be directly correlated with other predictive features, or outcome may be more related to stromal and epithelial components and associated genomic alterations in PDAC. Overall, 4 of the 6 tumour phenotypes were prognostically significant, and the markers, mutations, and microenvironmental cell types repeatedly selected across models had been linked to these phenotypes through prior analyses.

Copy number aberrations also proved to be unexpectedly informative predictors, though they require careful interpretation given that multiple drivers may coexist within each amplified or deleted locus. Poor prognosis was associated with *22q13.33* deletion (*HDAC10, MAPK11, MAPK12*), *6p21.1* amplification (*VEGFA, RUNX2, TFEB*), *10p11.21* amplification (*CCNY, CUL2, PARD3*), and *12q13.2* amplification (*ERBB3, PA2G4*). Conversely, improved outcomes were associated with *6p25.2* deletion (*PSMG2*), *3p21.31* deletion (*TREX1, PLXNB1, PFKFB4*), and *20p11.23* (*BFSP1, RRBP1*) deletion. Together, these findings highlight that broad tumour phenotypes, key stromal cell populations, and recurrent CNAs form the dominant axes of prognostic information, representing the core biological features most reliable for survival prediction.

## Discussion

Here we provide a panoramic map of immune, stromal, and epithelial cellular content in resected human PDAC, integrating proteomic, transcriptomic and genomic features. We observe a spectrum of tumour subtypes between classical and basal extremes with associated immune infiltrated and matrix-rich microenvironments that closely parallel the spectrum of patient outcome. Single-cell IMC charted 81 distinct cell types across the complex PDAC TME, from which tumour types and highly reproducible spatially defined microenvironments were identified revealing relationships between compartments consistent with interdependence or shared pathways. Supported by deep spatial proteomics and underlying genomics, our protein-guided phenotyping approach expanded upon transcriptional classifications to resolve previously unrecognized intermediate states. These tumours, which occupy the middle of the phenotypic spectrum, have historically been difficult to classify or target because they lack the clear transcriptomic or molecular hallmarks of either clearly classical or basal transcriptional subtypes, but are here identifiable by loss of lineage fidelity or mixed phenotype, as observed in poor-prognosis subtypes in other cancers^88,89^. This included a distinct hybrid phenotype expressing both canonical classical and basal markers, a low-ploidy S100A4+ mesenchymal phenotype, decreased or lost pancreatic epithelial transcription factor (edTF) expression and a CEACAM6 enriched phenotype. Considerable intra-tumour heterogeneity was observed and varied by subtype, but it was the presence of high-risk phenotypes and microenvironments within that heterogeneity that was associated with outcome, not the heterogeneity itself.

Basal epithelium and *ECM-rich* microenvironments, both linked to immune exclusion and poor outcomes, were more frequently associated with *KRAS* amplification than other poor-prognosis phenotypes. Given recent reports indicating that basal tumours tend to exhibit greater sensitivity to *KRAS* inhibition than classical subtypes^14,15^, we highlight that EMT-like phenotypes, such as S100A4+ classical-like tumours which appear less dependent on amplified *KRAS* signaling, are associated with resistance. This has important implications in anticipation of subtype-specific resistance to *KRAS* inhibitors and potential for synergy with immunotherapies and stroma targeting treatments. Despite immunotherapy failures in PDAC^90,91^, spatially resolved T cell activation was observed in many tumours providing hope for future therapeutic strategies. Accordingly, establishing a baseline measurement of cell content in treatment naïve PDAC is an important resource for comparison upon treatment. In untreated tumours, we identified *CD105+ fibrovascular* microenvironments, preferentially associated with classical subtypes and marked by increased vascularization, representing an intrinsic feature of classical PDAC. Notably, in our smaller number of treated patients, distinct *CD105+ CAF high* microenvironments persisted, suggesting selective preservation of this niche post-treatment, highlighting therapeutic vulnerabilities. Cross-omics analyses also refined our understanding of classical and intermediate tumour states, linking subtypes to pathway dysregulation and mechanisms of therapy resistance. Genomic aberrations in epigenetic regulators are associated with the absence of pancreatic edTFs and enrichment of dedifferentiated, poor-outcome tumour cell states, suggesting a connection to phenotypic plasticity. Altered TGF-β and interferon signaling were observed in poor outcome tumours and microenvironments associated with biophysical changes, while aberrant fucosylation, glycosylation and mucous production associated with gastric differentiation was observed across classical tumours. Further, we demonstrate spatially constrained cell type specific signaling patterns shaped by microenvironmental context especially within the many highly enriched myeloid cell states in pancreatic cancer.

Among all modalities tested, robust minimal biomarker sets derived from imaging features outperformed both clinical and genomic data in predicting overall survival. The strongest predictors combined tumour phenotypes, copy number aberrations, and specific CAF populations, while immune phenotyping—though not contributing directly to the models—showed correlations with selected features. Together, these results provide a high-resolution framework for understanding the spatial, molecular, and clinical heterogeneity of resectable PDAC and identify key phenotypes and microenvironmental features linked to patient prognosis.

## Acknowledgements

The authors are grateful to the Lunenfeld-Tanenbaum Research Institute (LTRI) at Mount Sinai Hospital for all technical support, the University Health Network (UHN) Cancer Biobank for specimen handling, and the Ontario Institute for Cancer Research (OICR) PanCuRx team for their dedicated partnership throughout this research. The authors acknowledge financial support from The Wallace McCain Centre for Pancreatic Cancer supported by the Princess Margaret Cancer Foundation for research programs involving patient consent and specimen acquisition, along with Canadian Institutes for Health Research (CIHR; PJT-175047), Canadian Cancer Society Research Institute (CCSRI) Breakthrough Team Grant (CCS 707708 Casper-PANC), Natural Sciences and Engineering Research Council (NSERC) Discovery Grant (RGPIN-2021-03404), Canada Foundation for Innovation (CFI), Government of Ontario – Ontario Research Fund (ORF) (40222 and 43479), OICR New Investigator Award (OICR IA-1-020), OICR Prosper-PANC (PanCuRx-FR2-SHS), and the Canada Research Chairs program (CRC-950-232945) to H.W.J. We wish to thank the Network Biology Collaborative Centre (NBCC) Proteomics Facility (RRID: SCR_025375) at the LTRI for mass spectrometry. The facility is supported by the CFI and the Ontario Government. N.S. acknowledges support from the Canadian Cancer Society (CCS) Research Training Award Program – Postdoctoral level in partnership with the Terry Fox Research Institute (TFRI) (CCS award 708064/TFRI grant # 708064) and Hold ‘Em for Life Oncology Fellowship from the Temerty Faculty of Medicine, University of Toronto. F.N. thanks CIHR for financial support in the form of the Canada Graduate Scholarships – Doctoral Program (CGS-D) award. S.D. is supported by a Canada Research Society (CRS) Doctoral Research Award and Ontario Graduate Scholarship (OGS). T.J.T. acknowledges support from the Agency for Science, Technology and Research (A*STAR) National Science Scholarship. We are deeply grateful to the patients for generously contributing tissue samples to this study. This work would not have been possible without their participation.

## Ethics declaration

H.W.J. has consulted for and received travel and research support from Standard BioTools unrelated to this work. K.R.C. reports consulting fees received from Abbvie Inc. and research funding from Sanofi and Standard BioTools, all unrelated to this work.

## Author contributions

N.S., F.N., S.D., T.J.T., E.S., J.L.G., G.M.O., and H.W.J. designed the study and reviewed the data. F.N., N.S., S.A., and H.W.J. performed IMC analysis and interpretation of data. N.S., F.N., S.D., E.S., E.L.C., G.A., J.L.G., H.W.J optimized and/or performed IMC. S.D., C.Y., M.J. and K.R.C. curated and performed analysis of publicly available datasets. N.S., S.D., F.N., E.S., C.J.W., B.S., Z.Y.L., J.P., A.C.G., and H.W.J. performed planning, experiments and analysis critical to interpretation of mass spectrometry. T.J.T., M.C., A.Z., S.C., J.M.W., R.C.G., F.M., S.G. acquired genomic sequencing and performed analysis. F.N. generated prediction model. A.B., R.G., S.B.L., M.M., A.D., B.G., J.K., R.C.G., F.N., G.M.O. and S.G. curated the clinical data and samples used within the tissue microarray. K.L. reviewed all samples and S.B.L. constructed the TMA. N.S., F.N., H.W.J. integrated all data and wrote the manuscript. All authors reviewed and approved the manuscript.

## Methodology

### Clinical Cohort

A total of 221 pancreatic ductal adenocarcinoma (PDAC) tumours were included in this study, obtained following surgical resection. The cohort comprised 195 treatment-naïve and 26 neoadjuvant-treated cases, with matched clinical data available for all 221 patients and whole-genome sequencing data available for 192 tumours. Tissue specimens were sourced primarily through the Ontario Institute for Cancer Research (OICR) PanCuRx initiative by affiliated surgeons from Toronto General Hospital (S. Gallinger). A subset of whole slide sections corresponding to select patients within the TMA were subsequently sectioned for laser capture microdissection (LCM). All samples were reviewed and verified by a board-certified pathologist (K.N.). From each tumour, four spatially distinct regions were sampled with a 1.25mm diameter coring needle and assembled into a tissue microarray (TMA), resulting in approximately 880 tissue cores, with cores from the same patient distributed across different slides. For spatial profiling, three 5-µm serial sections of the TMA were used to interrogate epithelial, immune and stromal compartments. All protocols for human sample collection and biobanking were approved by the appropriate institutional research ethics boards (Sinai 20-0170-E and 20-0178-E; UHN 20-5594 and 08-0767).

### Preparation and Sample Staining

No statistical method was used to predetermine sample size and all slides for each compartment were stained simultaneously. Formalin-fixed, paraffin-embedded (FFPE) sections were baked at 60°C for 1 hour, deparaffinized in xylene for 30 minutes, and rehydrated through a graded reagent alcohol series (100%, 100%, 96%, 90%, 80%, 70%, each for 5 minutes). Slides were rinsed in 1X Tris-buffered saline (TBS) for 10 minutes. Heat-induced epitope retrieval was performed in a 95°C decloaking chamber using Tris-EDTA buffer (pH 9.2) for 30 minutes. Following cooling, tissue sections were blocked for 1 hour at room temperature in buffer containing 3% BSA and 5% horse serum in 0.1% TBST. All cores per each slide were encircled with a hydrophobic barrier and incubated overnight at 4°C with a cocktail of individually titrated metal-tagged primary antibodies, as listed in Supplementary Table 2. For metal-conjugated secondary antibody use in panels, primary antibodies were incubated overnight at 4°C, followed by the secondary antibody for 1 hour at room temperature. Slides were washed three times with TBS and stained with iridium intercalator (500 nM) for 5 minutes for nuclear counterstaining. A final series of TBS washes was performed before slides were rinsed with deionized water and dried with pressurized air prior to IMC acquisition.

### Antibody Optimization and Conjugation

Antibodies were optimized on a panel of control tissues including human tonsil, spleen, appendix, and representative PDAC samples. Slide preparation followed the same protocol as described above, except for secondary antibody detection, primary antibodies were incubated overnight at 4°C, followed by the secondary antibody for 1 hour at room temperature. Slides were then washed three times with TBS and DAPI (2 μg/mL, 5 min) was used for nuclear counterstaining in place of the iridium intercalator. Specificity and co-localization were verified by immunofluorescence. Conjugation of validated antibodies to metal isotopes was performed using the Maxpar X8 Multi-Metal Antibody Labelling Kit (Standard BioTools) according to the manufacturer’s instructions and stored in Candor Antibody Stabilizer (Candor Biosciences) at 4°C.

### Image Acquisition

Image acquisition was conducted using the Hyperion and Hyperion XTi Imaging Systems (Standard BioTools). All acquisitions were performed at ∼1 µm resolution with laser ablation frequency of 400 or 800 Hz in a rasterized pattern using the Hyperion or Hyperion XTi, respectively. Acquisition order was randomized and blinded to patient identifiers and clinical metadata. Ablated tissue aerosol was ionized using inductively coupled plasma, and resulting isotopic ion reporters quantified using time-of-flight mass spectrometry to infer protein abundance. Raw data were compiled using Standard BioTools commercial acquisition software.

### Spillover Correction

To correct for isotopic spillover between channels for each panel, each individual metal-conjugated antibody was mixed with trypan blue and spotted onto agarose-coated slides. Quantification of isotopic contamination was performed by IMC. A spillover matrix was constructed and applied using the Bioconductor CATALYST R package^92^.

### Data Processing, Transformation, and Normalization

Images were first visually inspected using *Rakaia* (https://rakaia.io/) and converted to TIFF format. Single-cell segmentation was performed using *Mesmer* (DeepCell)^39^, implemented through our in-house analysis pipeline. Nuclear and membrane boundaries were defined using DNA, CD45, pan-cytokeratin, E-cadherin, and SMA signals to generate single-cell masks. Segmentation for the epithelial-focused panel was performed at a higher resolution (0.5 µm/pixel) compared to the immune and CAF panels (1 µm/pixel). For each region of interest (ROI), corresponding single-cell masks and expression matrices were generated. Marker intensities were capped at the 99.9th percentile to reduce the influence of extreme outliers and normalized to the maximum value observed for each marker across all samples.

### Cell Phenotyping and Microenvironment Annotation

Dimensionality reduction was first performed using principal component analysis (PCA), followed by clustering with PhenoGraph in R. Cell populations were annotated into defined epithelial subtypes, immune lineages, cancer-associated fibroblast (CAF) and endothelial states based on canonical marker expression. To enable spatial alignment of serial sections stained for epithelial, immune, and stromal compartments, we applied a computational overlay strategy. Ilastik object classification was performed on the panCK channel present in each serial section^93^. Affine transformation was then used to align the panCK objects across matched images of serial sections, and the learned transformation was applied to the x/y coordinates of the segmented cells. Finally, neighbor comparisons between compartments were used to verify alignment accuracy.

Epithelial lesions were defined as spatially clustered groups of ≥3 epithelial cells and matched across serial sections using the alignment strategy described above. Immune and stromal cells were assigned to their nearest lesion, and only microenvironments containing >3 cells with a corresponding lesion present in all three sections were retained, resulting in 9,890 high-quality microenvironments. PhenoGraph clustering of the cellular composition of these microenvironments identified eight distinct cross-compartment niches, each defined by recurrent combinations of endothelial, stromal, and immune cell types.

### Neighbourhood Analysis Using MILO

To assess differential spatial abundance of epithelial, immune, and stromal cell populations across clinical and genomic groups, we applied MILO, a statistical framework for differential abundance testing. MILO assigns cells to partially overlapping neighbourhoods on a k-nearest neighbour (k-NN) graph constructed from marker intensity profiles, and then tests each neighbourhood for significant enrichment between binary groups^47^.

Neighbourhoods were annotated as one of the predefined epithelial subtypes or aggregated immune and stromal compartments based on dominant cell label. For quality control, neighbourhoods with <70% label purity or with >50% of cells originating from a single patient were excluded. Differential abundance testing was then performed for three binary comparisons: ploidy (high, >2 average copy number, vs. low, ≤2), lymph node metastasis (positive vs. negative), and Moffitt RNA subtype (classical vs. basal). Results were visualized as beeswarm plots depicting the distribution of neighbourhood enrichments.

### *5-µm* IMC and Laser Capture Microdissection (LCM)

A consolidated 44-marker antibody panel targeting epithelial, immune, and stromal compartments was developed to enable identification of distinct tumour subtypes and microenvironments (Supplemental Table 4). This panel was applied on whole-tissue sections from select PDAC patient samples (Supplemental Table 5) for downstream laser capture microdissection and mass spectrometry (LCM-MS). Serial whole-slide sections were prepared at 4 µm thickness on standard charged slides for IMC, and at 10 µm thickness on polyethylene naphthalate (PEN) membrane slides for LCM of select microenvironment and epithelial regions. Staining of the consolidated panel using the IMC staining protocol described above was followed by acquisition on the Hyperion XTi imaging system (Standard BioTools) at 5 µm resolution in a rasterized pattern.

Following IMC acquisition, regions of interest were visualized and annotated using Rakaia. Visual alignment of tissue on FFPE PEN slides with annotated 5 μm image was performed. Using tissue landmarks and specific staining patterns on the 5 μm resolution image, regions of interest (ROIs) were freehand drawn in LCM software. ROIs were cut using an ultraviolet (UV) laser on the AccuLift Tissue Profiler (Targeted Bioscience). Cut regions were visually inspected with a microscope camera to orient, touch, and lift ROI off the slide with a 10 uL pipette tip. Cut tissue was placed into an 8-tube PCR strip containing 4 uL of 0.1% (w/v) n-dodecyl ß-D-maltoside (DDM) and used for protein extraction and downstream mass spectrometry-based protein quantification.

### Mass spectrometry (MS) of laser captured PDAC regions

Collected LCM regions were used for mass spectrometry analysis. Tissue lysates placed in 0.1% DDM (w/w) in 0.1 M ammonium bicarbonate (ABC) were heated at 95°C for 1h. Samples were reduced in 50 mM tris(2-carboxyethyl)phosphine (TCEP) in ABC and heated at 95°C for 30min. 1 μL of 60 mM iodoacetamide (in 0.1 M ABC) was added and incubated in the dark for 15mins. Proteins were digested in10 ng of Trypsin/Lys-C in 0.1M of ABC at 37°C for 16h. 1 μL of 10% formic acid was added to the samples to stop the digestion and samples were brought to a final volume of 20 μL with HPLC water.

For data-independent acquisition (DIA) of LC-MS/MS, all of the digested peptides were analyzed using a nano-HPLC (high-performance liquid chromatography) coupled to mass spectrometry (MS). Peptides were separated on an IonOpticks Aurora3 Elite column (cat. AUR3-15075C18) using the 40SPD Whisper Zoom method, with 50°C column temperature analysed on a timsTOF SCP mass spectrometer. MS acquisition was performed in PASEF mode across a mass range of 100–1700Da. Singly charged ions were excluded using a polygonal filter. Peptide identification and quantification were performed using Spectronaut v19 (Biognosys) with the directDIA+ workflow and the Spectronaut-generated Human_PDB_2023 library (Supplemental Table 7). Default search parameters were used, with no signal imputation or cross-run normalization.

### scRNA-seq

8 publicly available single cell (sc) RNA-seq datasets from human primary PDAC (n = 163) were used to project associated genes from the Spectronaut protein pivot table onto single-cell transcriptomic cell types. Following initial sample filtering, all datasets provided in CellRanger output format underwent in-house preprocessing and quality control. Doublet detection was performed using doubletFinder_v3 from the DoubletFinder v2.0.3 package^94^, and identified doublets were excluded from downstream analysis. Low-quality cells were removed based on standard thresholds: cells were filtered out if they exhibited >20% mitochondrial RNA content, expressed fewer than 500 genes, or contained fewer than 1,000 total transcripts. Remaining UMI counts were normalized using a per-cell size factor and subsequently log2-transformed with the logNormCounts function from scuttle v1.8.4^95^, and unbiased cell type recognition was performed at the single cell level using SingleR^96^ from SingleR v.2.0.0, as previously described(*1*).

Single cells annotated by SingleR were categorized into five groups - immune, endothelial, cancer-associated fibroblasts (CAFs) and mural cells, ductal cells, and acinar and B cells. For each group, the raw count data was normalized using a per-cell size factor (computeSumFactors from scran v1.34)^97^ and subsequently log2-transformed with the scater v1.34 logNormCounts^95^ function. To remove any remaining potential doublets, cells annotated as immune, endothelial, or cancer-associated fibroblasts (CAFs) that expressed epithelial marker genes (KRT19, KRT5, and EPCAM), as well as ductal cells lacking expression of these markers, were excluded from downstream analyses. For each of the five groups, UMAPs were generated using umap v0.2.10.0 on batch corrected data derived from batchelor’s fastMNN (v1.22)^98^. Leiden clustering^99^ (Scanpy v1.0.2) was applied to each group, and differential expression on normalized, non-batch-corrected data was used to identify marker genes for each cluster. Clusters were manually annotated based on known markers, and differential expression results informed IMC panel design.

### Mass Spectrometry and scRNAseq Integration

A total of 275 laser capture microdissected (LCM) regions were annotated based on 5 μm IMC-defined microenvironments. Differential protein expression across environments was evaluated using the Wilcoxon rank-sum test via the wilcox.test function in base R version 4.4.3, applied to data from the Spectronaut protein pivot table. Filtered values in the protein matrix were either imputed as zero or, for comparison, to the lowest 10% of values detected within the corresponding sample. To avoid excluding proteins shared across similar microenvironments, the environments were grouped as follows: (CH, CL, CN, B), (M, I, IS), and (S, E, CCF, CM), with each microenvironment compared against all samples outside its group. Gene–environment pairs were retained only if the mean expression of the gene within the microenvironment reached at least 75% of the maximum mean expression observed across all environments. Then, the genes corresponding to the 937 differentially expressed proteins were plotted in single-cell RNA sequencing (scRNA-seq) data and genes which were highly variable in the scRNA-seq data were selected manually for visualization. The highly variable genes for each microenvironment were entered into gprofiler using the default library combined with a library generated from curated PDAC genes, to determine significantly enriched pathways in each of the microenvironments.

### Whole-Genome Sequencing (WGS)

Whole-genome sequencing data from TMA-matched samples (n = 192) were used to identify copy number alterations and mutations in the same patient cohort, providing additional context to the functional and spatial features of the microenvironments. Each donor underwent multi-regional laser capture microdissection followed by whole genome sequencing as previously described^100^. A given donor was encoded positive or negative for a mutated gene or a non-synonymous alteration. For somatic copy number alterations (CNA), we employed the use of GISTIC2.0 to identify recurrently broad and focal genome alterations to derive a set of genomic regions for association^101^. The resultant peak level copy number values per tumour were directly used for associations. Both mutations and CNAs were tested for association with a given tumour phenotype utilizing a generalized linear model, taking the logit-transformed proportion as the dependent variable. For gene level mutations, only genes with mutations observed in 10 or more patients were used. Each model was adjusted for average overall ploidy, cellularity and total cell counts, represented as a fractional rank-transformed covariate. Adjustments for multiple testing were made utilizing the Benjamini-Hochberg approach.

### Machine Learning Prediction

The primary outcome for prediction was long-term survival (>2 years, n = 104) versus early mortality (≤2 years, n = 93) after diagnosis. Survival time was discretized to account for outlier patients with >8 years survival (n = 10) and to address right-censored continuous overall survival data. Patients alive with <2 years of follow-up data were excluded (n = 4). All machine learning analyses were conducted in Python (v3.9) using the Stabl framework^87^, which implements nested cross-validation with feature stability selection. The outer loop consisted of 100 stratified random splits (80% training, 20% testing) with a fixed random seed for reproducibility. Within each outer split, model hyperparameters were tuned using a five-fold cross-validation repeated five times (25 inner resamples).

Six independent data modalities were used in the initial analyses: single-nucleotide variants (SNVs), copy number aberrations (CNAs), clinical variables, protein marker intensities, spatially normalized cell type densities, and microenvironment proportions. Preprocessing was performed separately for each data type. SNVs were restricted to genes mutated in at least 10 patients and filtered to exclude very large genes (>2000 amino acids) whose mutation frequency correlated with overall tumour mutational burden, with the exception of large genes known to play a role in DNA damage repair or mutation rates (ATM, KMT2A, KMT2D). CNAs, protein marker intensities, cell type densities, and microenvironmental features were standardized (mean = 0, SD = 1). Clinical variables were either dummy-encoded (categorical) or z-scored (continuous). Within each inner loop, Stabl generated artificial features matched to each dataset’s distribution to control for false discovery. Feature selection was performed using 150 bootstrap subsamples; features whose selection frequency exceeded the empirical false discovery threshold (FDP+) were retained. Selected features were used to train Lasso models (logistic regression with L1 penalty), with the area under the ROC curve (ROC-AUC) as the optimization metric. Models trained on the selected features were then evaluated on the 20% held-out test set.

After completing feature selection independently for each data modality, features that appeared in at least 50% of iterations were merged across data types into a single integrated feature matrix. The fused dataset was re-analyzed with the Stabl workflow to identify a refined set of robust predictive features distinguishing patients with long-term survival (>2 years) versus early mortality (≤2 years) after diagnosis.

---

**Extended Fig1:**
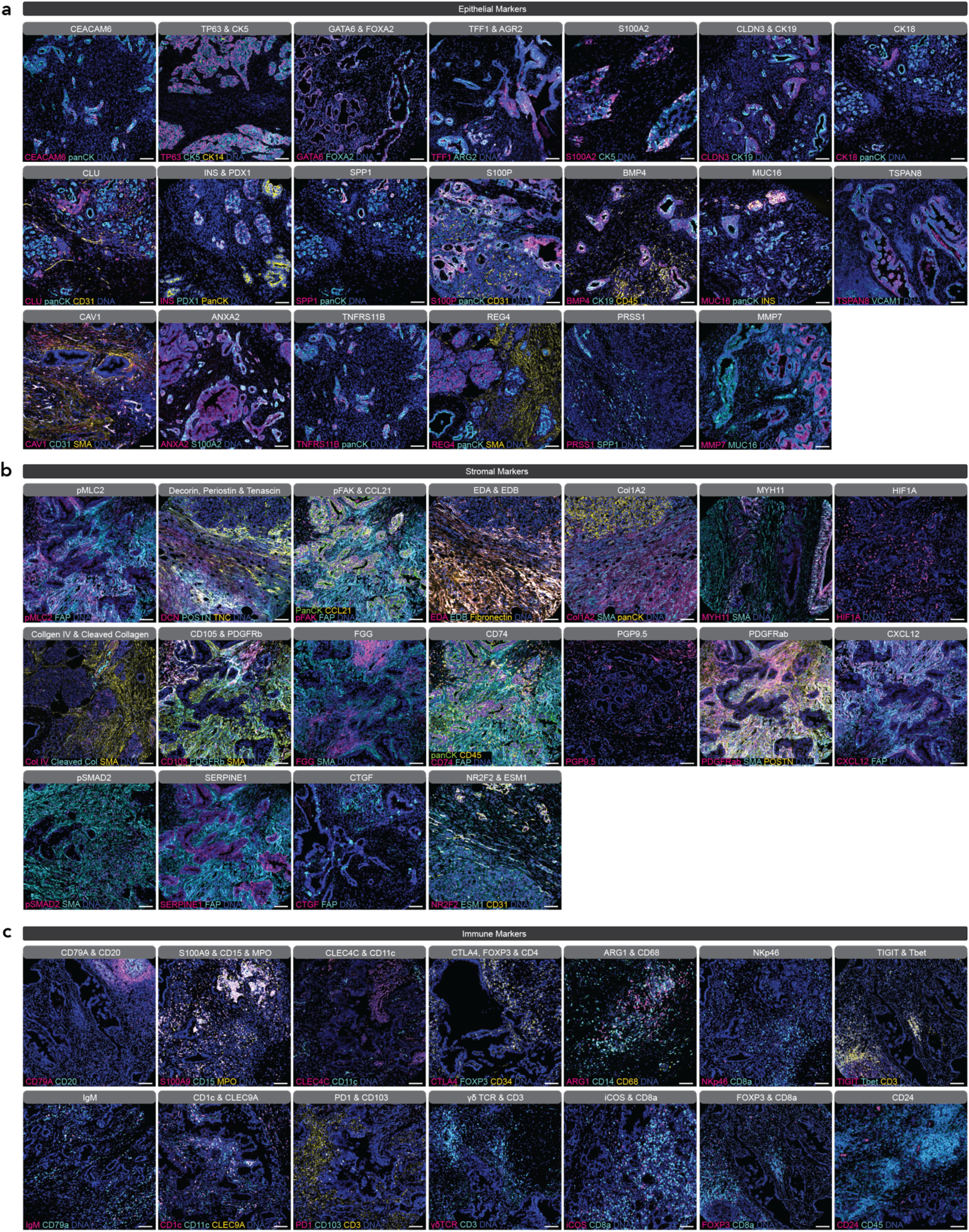
Validation of previously unpublished antibodies for multiplex IMC imaging targeting (a) epithelial, (b) stromal, and (c) immune compartments. Scale bar represents 50 μm. The full resolution images will appear in final publication.

**Extended Fig 2:**
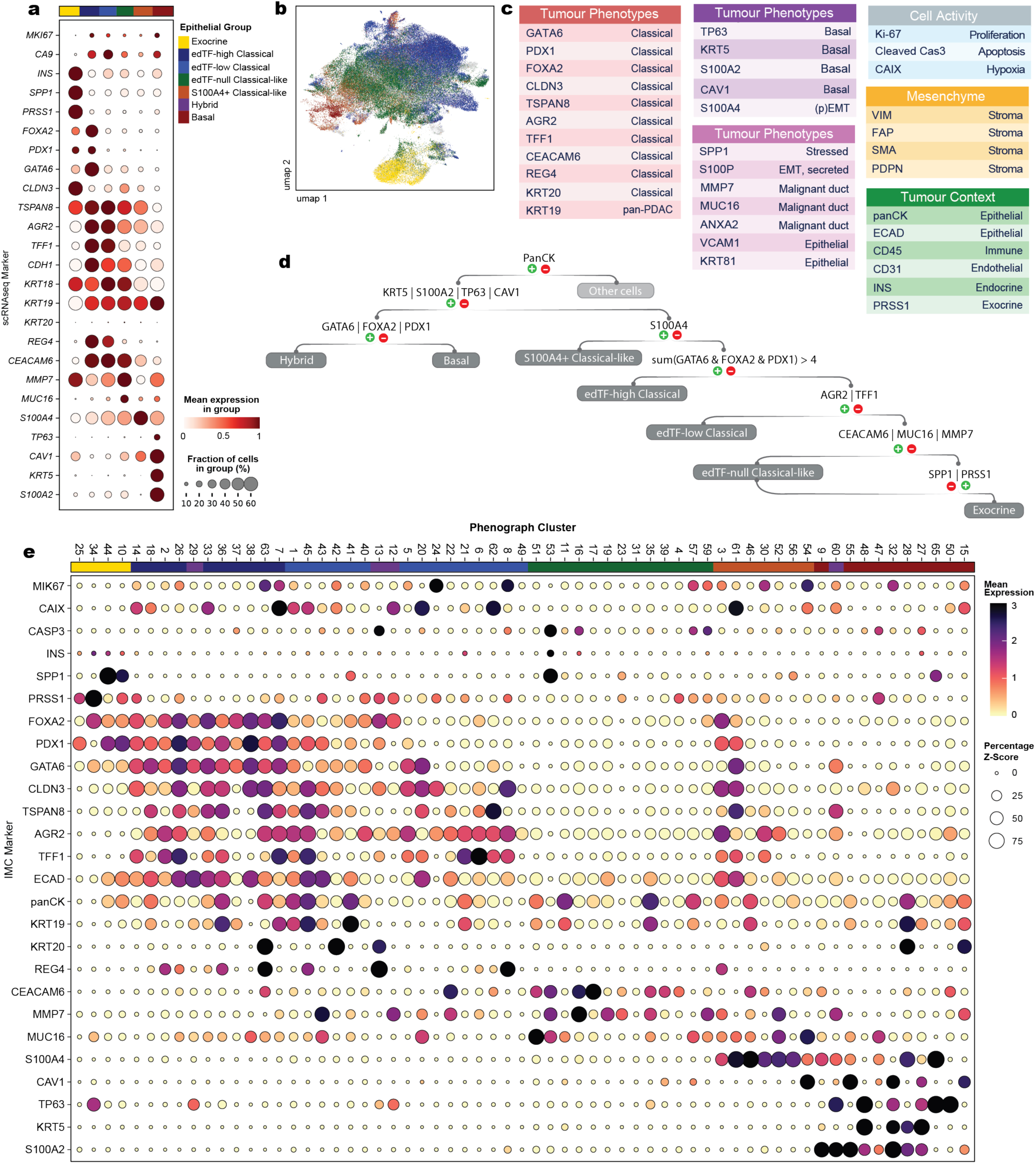
Epithelial IMC panel resolves 7 distinct phenotypes validated by transcriptomic expression. **a**, Average expression of each epithelial panel marker in scRNAseq data when cells are gated as shown in UMAP (b). scRNAseq, single cell RNA sequencing. **b**, UMAP of merged scRNAseq from 8 datasets (n = 163 patients), with areas representative of the 7 IMC-based phenotypes highlighted. **c**, Epithelial antibody panel design for IMC covering markers for tumour phenotypes, cell activity, mesenchyme and tumour context. IMC, imaging mass cytometry. **d**, Decision tree used to annotate the original unsupervised clustering of 66 epithelial clusters. **e**, Heatmap of marker expression across all epithelial clusters, with the assigned phenotype annotations shown in the top colour track.

**Extended Fig 3:**
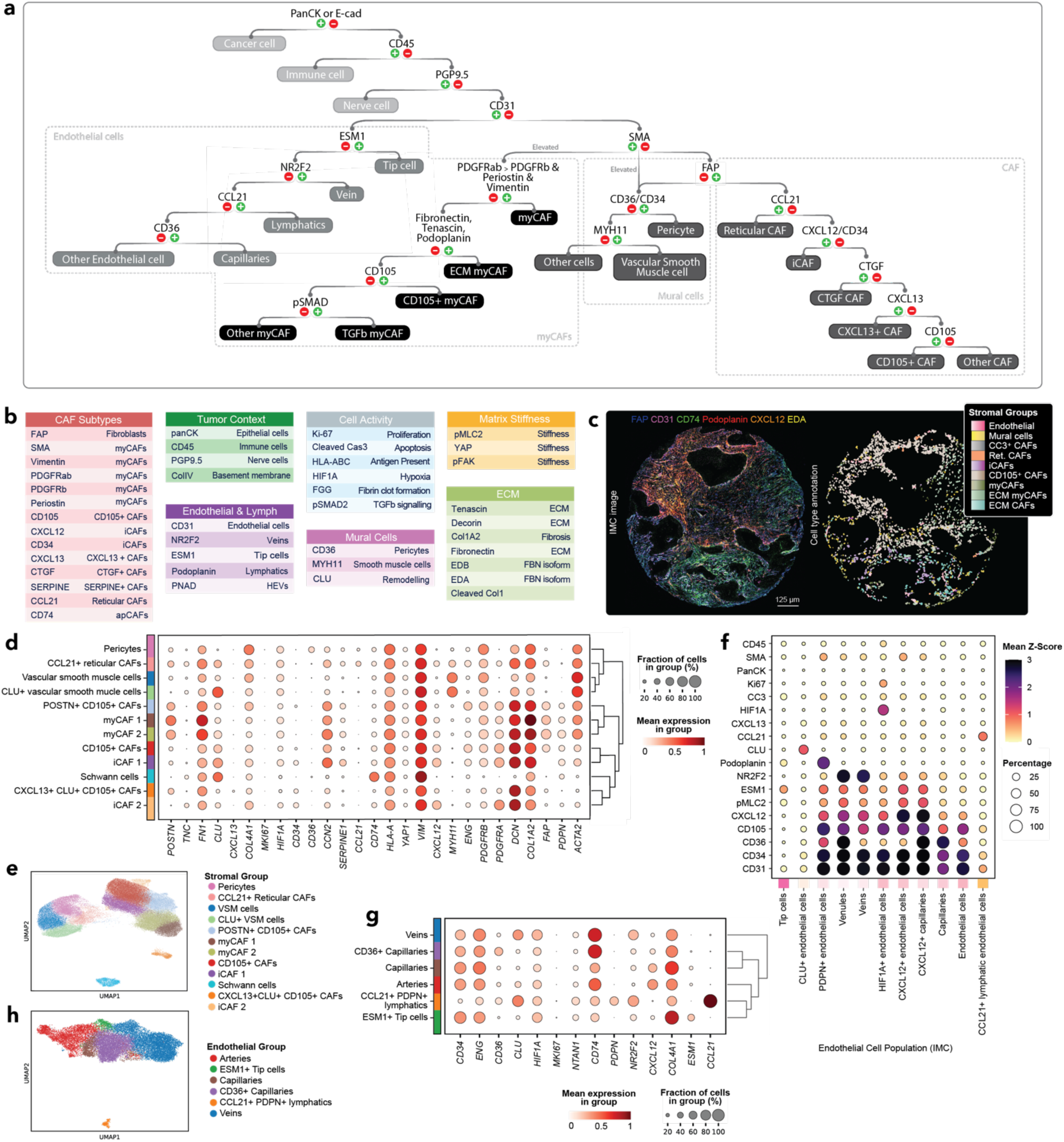
PDAC stromal panel validation and fibroblast/endothelial cell mapping using scRNAseq. **A**, Schematic flow chart of stromal, endothelial, and mural lineage assignment based on marker selection for the stromal panel. Endothelial subtypes not shown as standalone populations are annotated by their dominant marker, **b**, Completed list of stromal and endothelial markers included in the IMC panel, categorized by CAF subtypes, tumour context, endothelial cells and lymphatics, cell activity, mural cells, matrix stiffness and ECM. IMC; imaging mass cytometry; ECM, extracellular matrix. **c**, Representative IMC image (*left*) of select stromal markers and corresponding cell type annotations (*right*) generated via *cytomapper.* **d**, Bupple plot displaying expression of IMC panel markers across fibroblasts in sing cell RNA sequencing (scRNAseq) data from PDAC tumours (n = 163 patients), illustrating marker specificity, **e**, UMAp of annotated fibroblast populations derived from scRNAseq, corresponding to the bubble plot in (d). **f**, Average IMC expression of markers across endothelial cell types from PDAC samples stained using the stromal panel of antibodies. **g**, Bubble plot displaying expression of IMC panel markers across endothelial populations from scRNAseq data (n = 163 patients). **h**, UMAP of annotated endothelial cell types corresponding to bubble plot in (g).

**Extended Fig 4:**
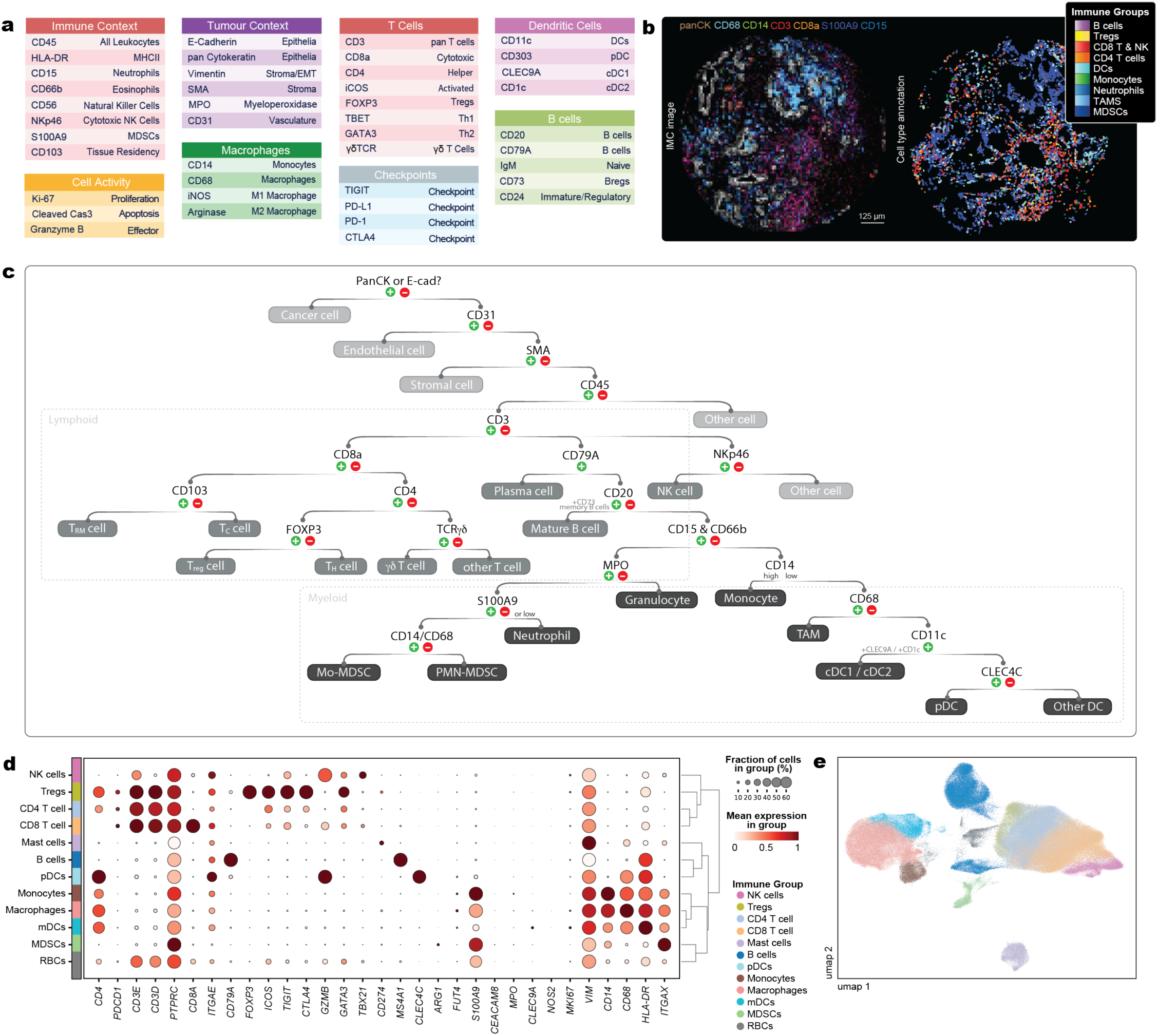
PDAC immune panel validation and hematopoetic cell lineage mapping using scRNAseq. **a**, Complete list of immune markers included in the IMC panel, categorized by immune context, tumour context, T cells, dendritic cells, B cells, macrophages, checkpoints, and cell activity. IMC; imaging mass cytometry. b, Representative IMC image *(left)* of select immune markers and corresponding cell type annotations *(right)* as generated via *cytomapper.* c, Schematic flow chart of lineage assignment based on markers in this study. d, Bubble plot displaying expression of IMC panel markers across hematopoetic cell types in single cell RNA sequencing (scRNAseq) data from PDAC tumours (n = 163 patients), demonstrating marker specificity. e, UMAP visualization of annotated cell populations derived from scRNAseq, corresponding to the bubble plot in (d).

**Extended Fig 5:**
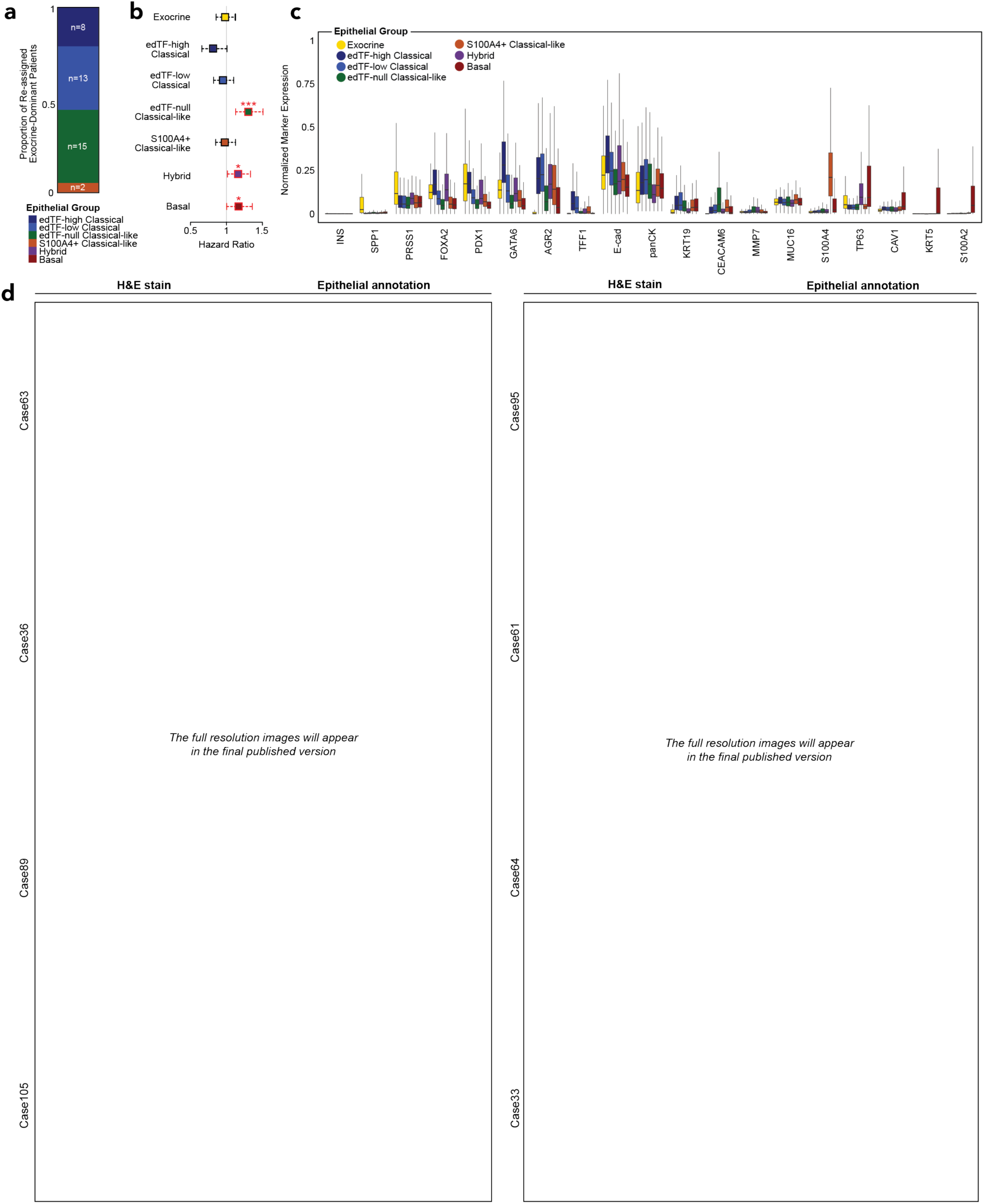
Protein expression profiles, histopathology (H&E), and single-cell segmentation of 7 epithelial phenotypes in PDAC tumours. **a**, Stacked bar displaying the proportion of exocrine-dominant patients re-assigned to each tumour subtype, with corresponding patient counts indicated. **b**, Hazard ratios (left) for each *epithelial* tumour type, calculated using Cox proportional hazards regression. Points outlined in red are statistically signficant. Asterisks above each point indicate significance level: Basal: P=0.044(*), Hybrid: P=0.031(*), edTF-null Classical-like: P<0.00036 (***). **c**, Boxplots showing marker expression distribution, normalized 0-1, per annotated epithelial group in single cell data. Outliers omitted. **d**, Matched serial sections of H&E and segmented IMC images visualized using the *cytomapper* R package, where each epithelial group represented at least once. From left to right, top to bottom, the cores shown have the IDs PDAC1_Sector3Row1Column8_Punch1_Case63_ID_1_93605 (edTF-high Classical), PDAC1_Sector4Row1Column8_Punch1_Case95_ID_G_31568 (Exocrine), PDAC1_Sector2Row2Column4_Punch1_Case36_ID_1_69515 (Hybrid), PDAC1_Sector3Row1Column6_Punch1_Case61_ID_1_92541 (Exocrine), PDAC1_Sector4Row1Column2_Punch1_Case89_ID_G_26400 (S100A4+ Classical-like), PDAC1_Sector3Row2Column1_Punch1_Case64_ID_1_93724 (Basal), PDAC1_Sector4Row3Column3_Punch1_Case105_ID_G_9208 (edTF-null Classical-like), PDAC1_Sector2Row2Column1_Punch1_Case33_ID_1_67113 (edTF-low Classical).

**Extended Fig 6:**
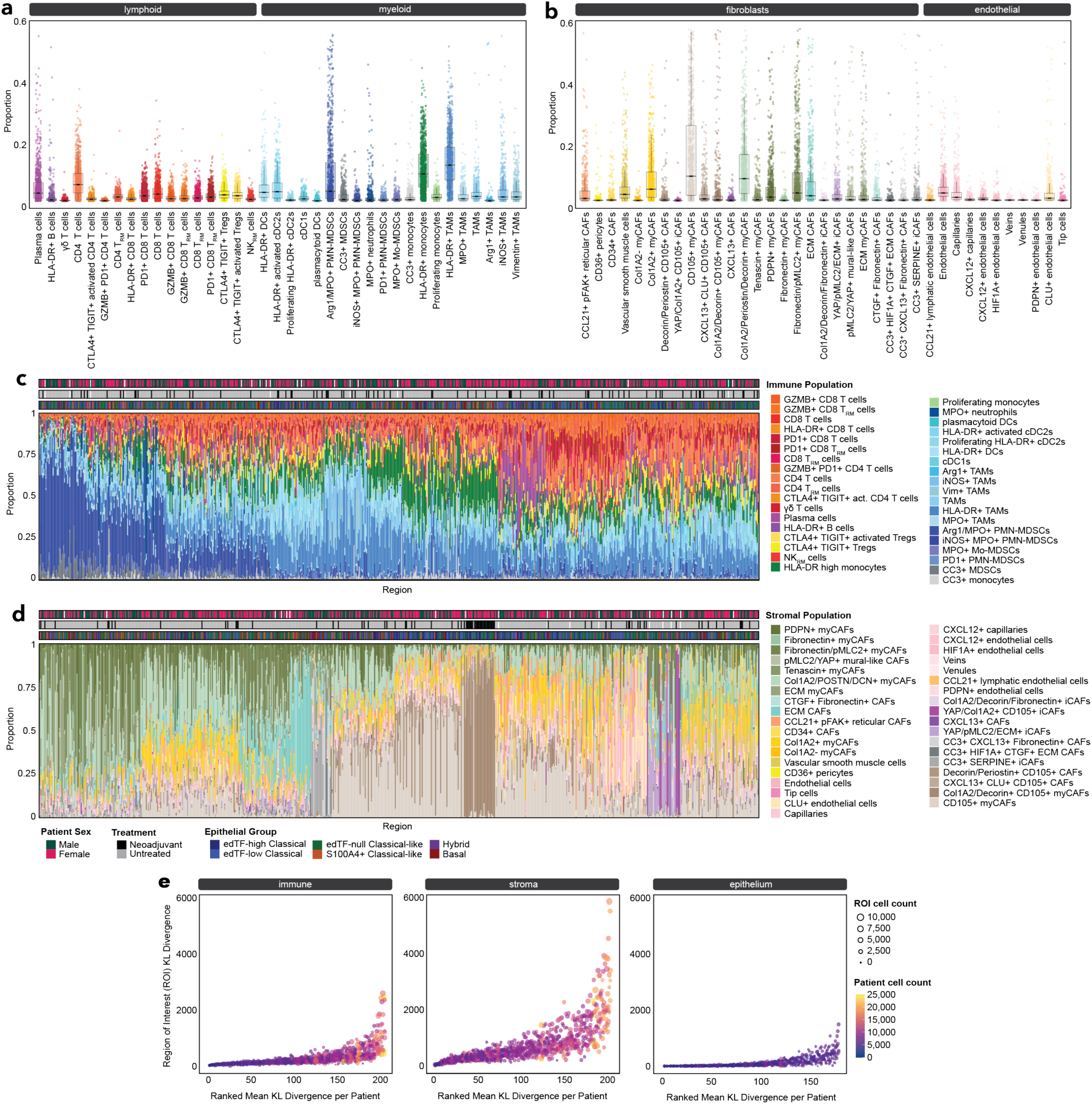
Characterization of immune and stromal proportions and regional heterogeneity in PDAC tumours. **a-b**, Boxplots showing the distribution of (a) immune and (b) stromal (fibroblast and endothelial) cell types as a proportion of total cells within their respective compartments Each dot represents a ROI, with values corresponding to the proportion of the specified cell type relative to all cells in that compartment. ROI, region of interest. **c-d,** Stacked bar plot displaying the proportional composition of (c) immune and (d) stromal cell types within each region of interest (ROI; n = 823). Each bar represents a single ROI, with the y-axis indicating the relative abundance of each immune or stromal cell population. Track legends above stacked bar plots display patient sex, treatment history, and the dominant epithelial phenotype associated with the patient from which each ROI is derived, as denoted below panel d. e, Quantification of immune, stromal or epithelial heterogeneity across patients using KL divergence. Each dot represents a ROI plotted by its KL divergence from the patient’s mean immune, stroma or epithelial composition. X-axis shows patients ranked by their average KL divergence across ROls. Higher KL divergence values indicate greater intra-patient spatial heterogeneity in respective compartment’s cell composition. KL, Kullback Leibler.

**Extended Fig. 7:**
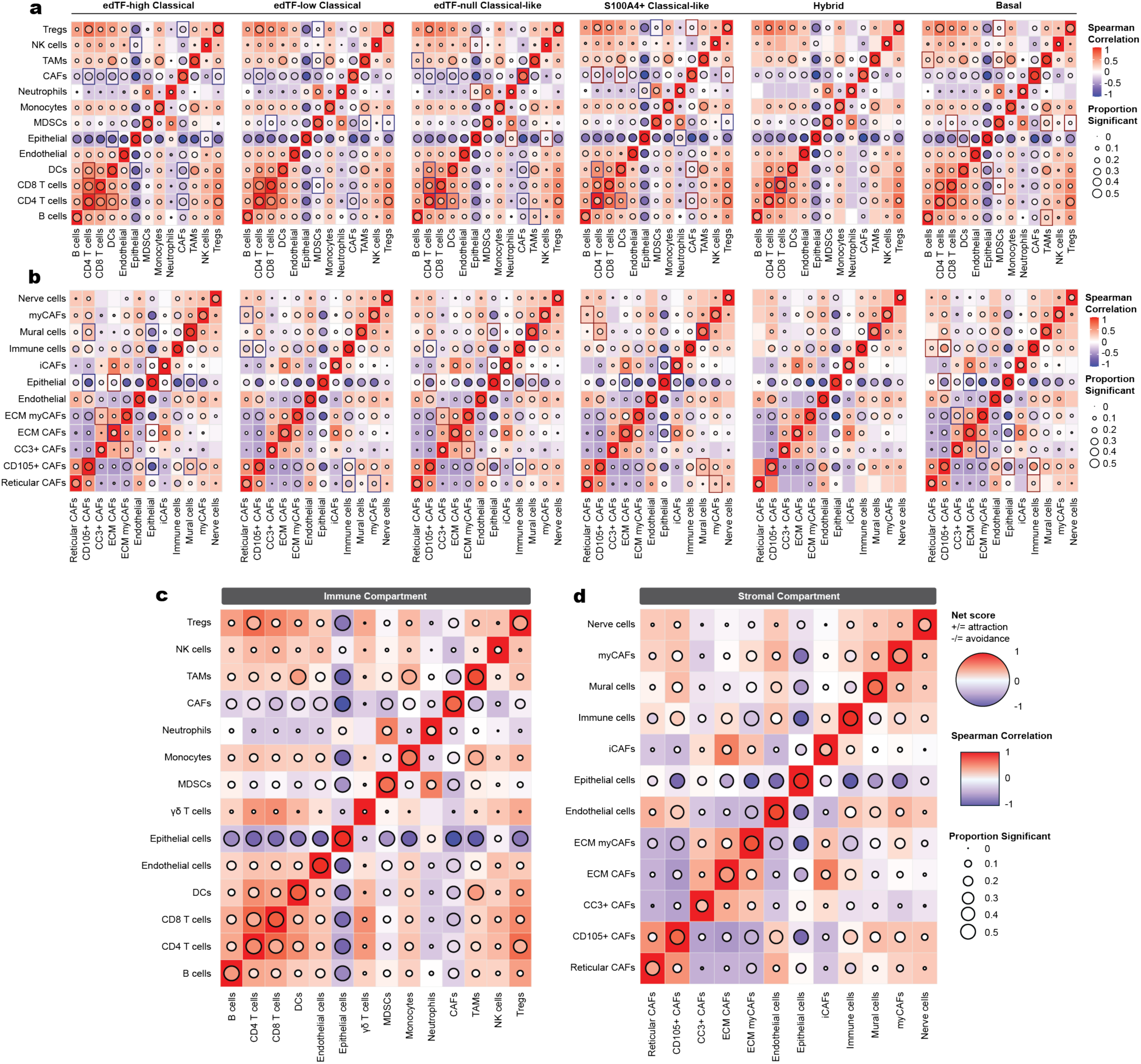
Pairwise spatial interactions across immune and stromal compartments. **a-b**, Spearman correlation heatmaps displaying how often pairs of cell types co-occur within individual patient images, stratified by dominant tumour subtype (labelled at the top). The tile colour represents the strength of the correlation, while circle colour and size indicate the avoidance/interaction strength and direction between immune cells (a) or stromal cells (b). The tile outline denotes statistically significant differences in interaction patterns between the dominant tumour type that interaction belongs to and at least one other tumour type. Each analysis includes epithelial, endothelial and stromal or immune cells depending on the panel. **c-d,** Overall Spearman correlation heatmaps across all immune populations (c) and stromal populations (d), summarizing co-occurrence and interaction trends across the full cohort without stratifying by dominant tumour subtype.

**Extended Fig 8:**
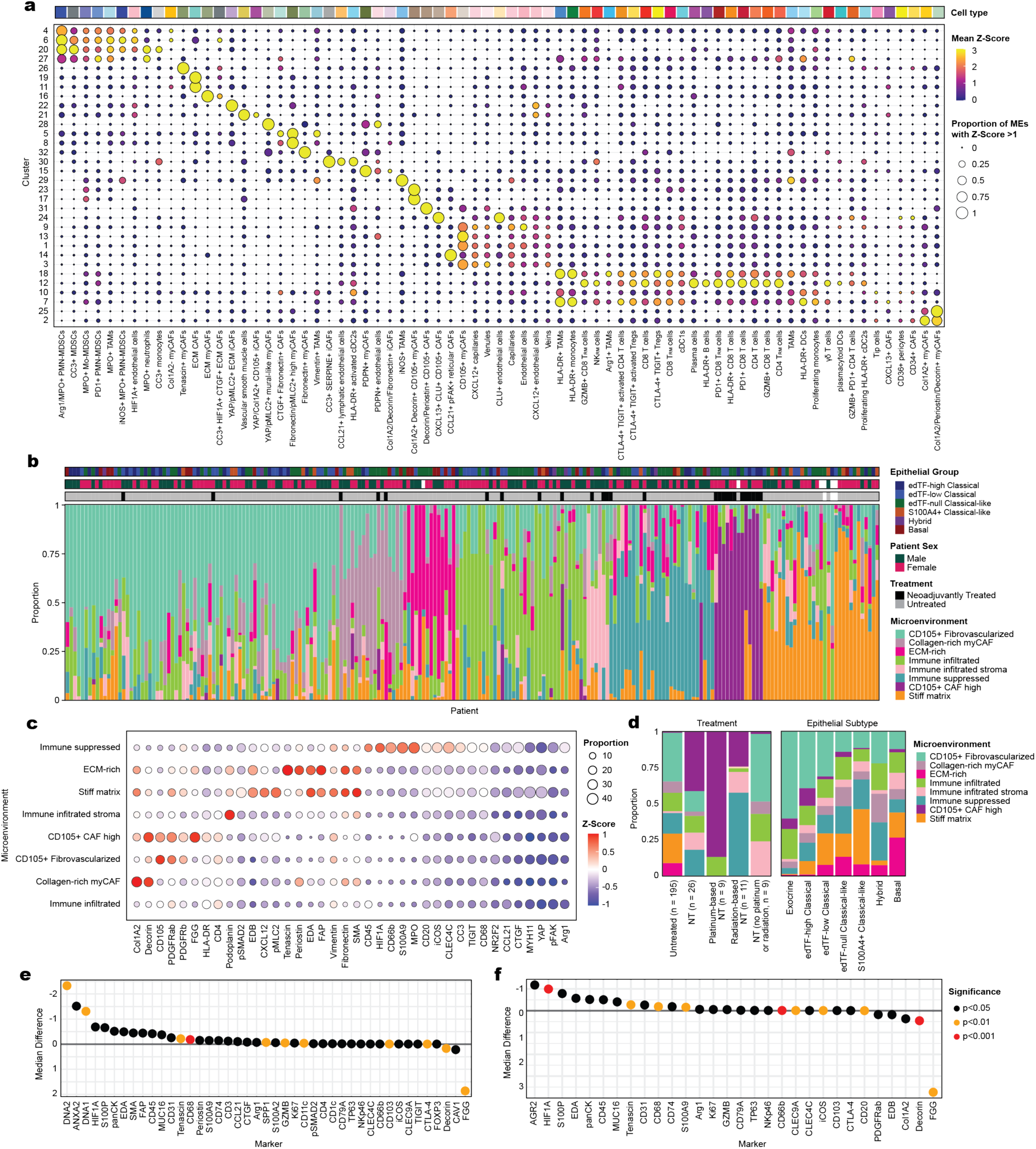
PDAC microenvironments display distinct IMC-defined phenotypes and treatment-associated remodeling across spatial compartments. **a**, Bubble plot displaying mean expression of each cell type per microenvironment cluster. Circle size represents the proportion of cells that express each corresponding marker above a Z-score of 1. Markers were Z-scored and the colour represents the average Z-score for each marker per cell type. Colour track represents each cell type. ME, microenvironment. **b**, Stacked bar plot displaying the proportion of each of the eight microenvironments per patient. Top annotations indicate the dominant epithelial subtype within that patient, patient sex, and treatment status (neoadjuvant-treated or treatment naive). **c**, Bubble plot depicting primarily functional marker expression within each microenvironment. Here, *’microenvironmenf* denotes immune and stromal cell expression within these microenvironments and excluded expression on epithelial lesions. Circle size represents the proportion of cells that express each corresponding marker above a Z-score of 1. Markers were Z-scored and the colour represents the average Z-score for each marker per cell type. Plot includes select immune and stromal markers along with functional markers. **d**, Stacked bar plot representing the proportion of microenvironments from patients comprising each treatment group *(left)* and epithelial subtype *(right).* NT, neoadjuvant treatment. **e-f**, Mean difference plots showing the average marker expression differences between (d) all neoadjuvant-treated patients (n = 26) vs untreated patients (n = 195) and (e) platinum-based neoadjuvant-treated patients vs all others. Markers to the right are enriched post treatment, while those to the left and reduced. Dot colour reflects significance (p < 0.05, 0.01, 0.001).

**Extended Fig 9:**
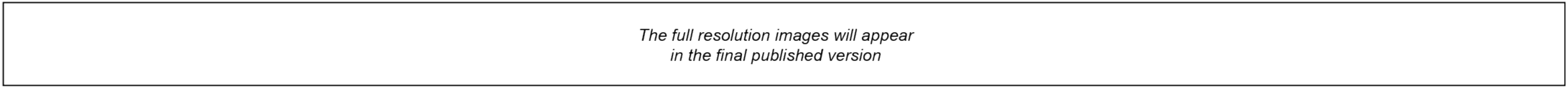
Representative 5-µm resolution IMC whole-slide images. These 42-channel multiplexed images were used to annotate tumour phenotypes and microenvironments corresponding to the patterns described in the 1-µm resolution IMC images of tissue microarray cores. The annotated images then guided laser-capture microdissection of serial whole-slide sections for mass spectrometry measurements.

**Extended Fig 10:**
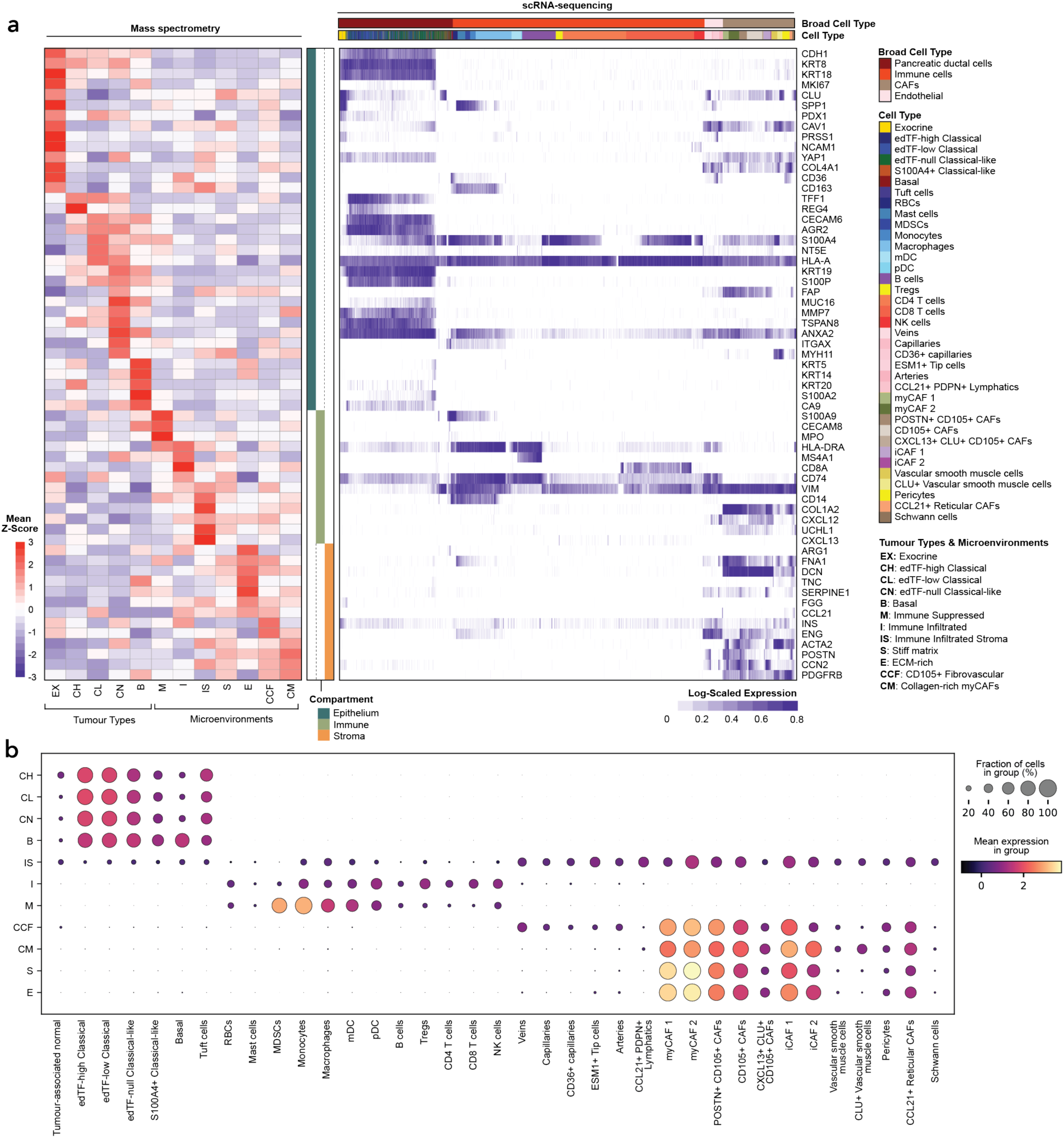
Spatially resolved epithelial subtypes and microenvironments exhibit overlap with IMC expression across multi-omic data. **a**, *Left* Heatmap of average protein abundance across LCM regions profiled by MS (LCM-MS), corresponding to the microenvironments annotated in (a). Columns represent individual microenvironments; rows represent proteins within our imaging mass cytometry panels, z-score normalized across all regions. LCM, laser capture microdissection; MS, mass spectrometry; IMC, imaging mass cytometry. *Right* Expression of genes encoding the proteins identified by LCM-MS, as detected in single-cell RNA-seq data and stratified by IMC-defined microenvironments. Columns represent individual cells, annotated using SingleR based on known marker genes. Gene expression is log-normalized and row-scaled to the maximum expression per gene. ECM, extracellular matrix; CAFs, cancer-associated fibroblasts. **b**, Bubble plot displaying scRNAseq-annotated cell populations along the x-axis and the micronevironments and epithelial subtypes they are associated with on the y-axis. Bubble size reflects the abundance or frequency of each cell population within the corresponding context.

**Extended Fig 11:**
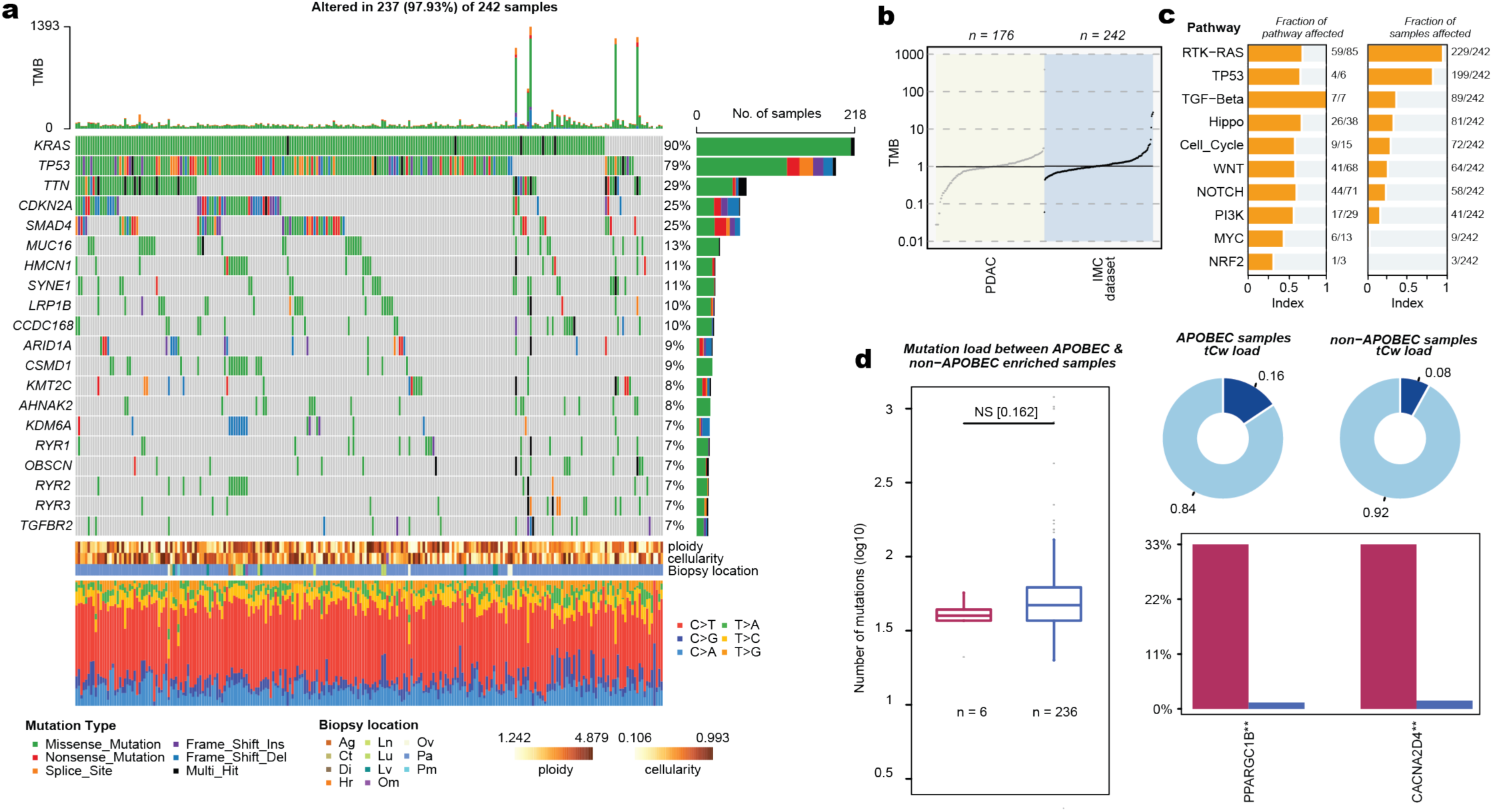
Overview of whole-genome sequencing in PDAC tumours. **a**, Oncoprint displaying the most frequently altered genes in PDAC across 242 tumour samples (192 unique patients), including mutations (missense, nonsense, frameshifl, splice site) and complex events. Bars on the right indicate the percentage of samples affected per gene; top barplot shows the number of mutations per sample. Ag, adrenal gland; Ct, cutaneous; Di, diaphragm; Hr, heart, Ln, lymph node; Lu, lung; Lv, liver; Om, omentum; Ov, ovary; Pa, pancreas; Pm, peritoneum. b, TMB displayed on log10 axis; each dot represents a patient, with the black horizontal line denoting the median TMB (1.02 mutations/MB). Comparison of TMB against TCGA-PAAD Cohort. TMB, tumour mutational burden. c, Fraction of signalling pathways affected by mutations. d, Comparison of number of mutations (log10 scale) between APOBEC and non-APOBEC enriched samples shows no enrichment of these signatures in our cohort.

**Extended Fig 12:**
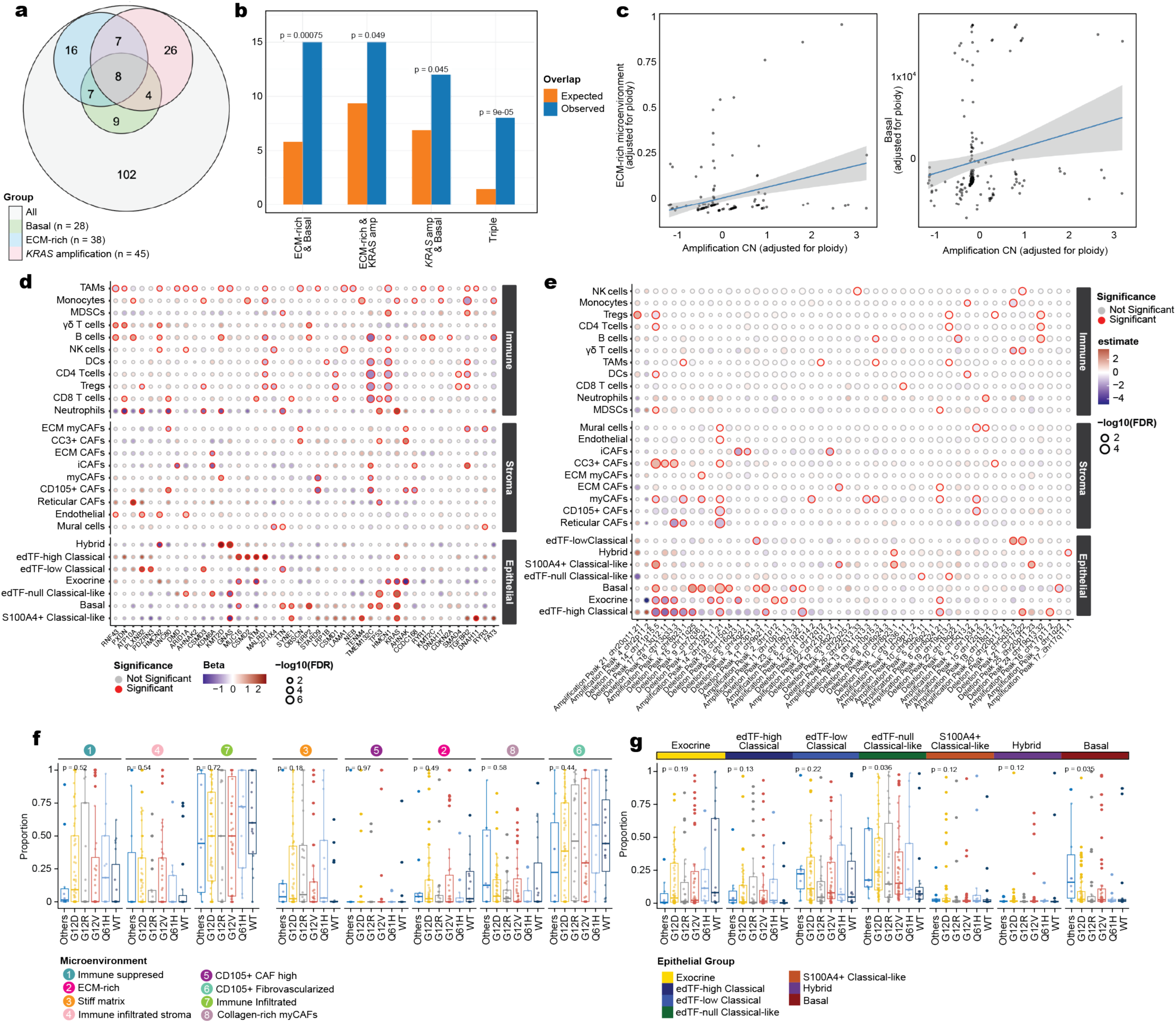
Subset WGS analysis and cell type-specific SNP and CNA associations. **a**, Venn diagram illustrating the number of patients with matched whole genome sequencing (all, n = 182 patients) categorized into three distinct groups: basal (patients with >20% of their tumour cells annotated as basal; n = 28), ECM-rich (presence or absence of ECM-rich microenvironments; n = 38), and *KRAS* amplification (patients with a copy number >duploid at the *KRAS* locus; n = 45). **b,** Bar plot showing the expected number of patients overlapping between basal cell presence, ECM-rich microenvironments and *KRAS* amplification groups based on 1e+05 simulations using the observed frequencies of each feature across the cohort (n = 182). **c,** Correlation between *KRAS* amplification and either the presence of ECM-rich microenvironments (left) or basal cell content (right), adjusted for ploidy. **d-e,** Bubble plots representing associations between cell types (immune, stromal and epithelial) and (d) SNPs or (e) CNAamplification and deletion identified using GISTIC2.0, summarized to their respective cytogenetic bands. Statistically significant associations are highlighted in red. SNP, single nucleotide polymorphism. **CNA,** copy number alteration. **f-g,** Box plots representing associations with specific *KRAS* mutations utilizing Kruskal-Willis test for microenvironments (f) and epithelial groups (g). *WT,* wild-type.

